# What has changed in 20 years? Structure and function of soft-sediment macrofauna in a subarctic embayment, Newfoundland (Canada)

**DOI:** 10.1101/2023.10.29.564610

**Authors:** Ivana Komendić, Bárbara de Moura Neves, Patricia A. Ramey-Balci

## Abstract

Understanding how natural and anthropogenic disturbances affect the structure and functioning of marine ecosystems and their associated services is central to predicting future dynamics. Placentia Bay is an Ecologically and Biologically Significant Area (EBSA) in the north Atlantic exposed to multiple stressors. To investigate changes in the community and functional structure of soft-sediment macrofauna as well as environmental drivers of observed variation, we compared contemporary (2019–2020) and historical (1998) samples at eight stations (*n*=77) collected 21 years apart. Although community and functional structure significantly differed between these two time points, functional traits were maintained (i.e., no loss of 36 trait modalities). Thirty-seven percent of species/taxa where only observed in either the historical or contemporary community, and the contemporary community exhibited a lower density of macrofauna but had similar richness, resulting in higher evenness and diversity. Highly tolerant subsurface deposit feeders having small body sizes (<10mm) and direct development dominated the historical community, whereas the contemporary community had nearly equal proportions of surface and subsurface deposit feeders with small to medium body sizes (<10–50 mm) with pelagic larvae, and the proportion of highly tolerant species/taxa was reduced. These changes likely reflect the large reduction in polychaetes (91 vs. 58%) and increased bivalves (4 vs. 25%) relative to the historical time point. Community variation was driven by changes in the sedimentary habitat. Contemporary versus historical sediments were ∼4.5x coarser and had higher levels of sedimentary organic matter. This work contributes to advancing our understanding of relationships between benthic macrofauna, functional traits, and the sedimentary habitat in coastal environments.

## Introduction

With global climate change altering ecosystem conditions, information pertaining to both community structure and ecosystem function are necessary to understand and determine important drivers of environmental change and predict the resilience of communities (Biggs et al. 2020; Vause et al. 2019). Like many coastal environments our study site in Placentia Bay Newfoundland, located in the Northwest Atlantic, is exposed to multiple stressors including rising sea surface temperatures, storm events, tanker traffic, and aquaculture. In the marine environment, soft-sediment benthic macrofauna (herein defined as organisms retained on a 500 µm screen) can exhibit high diversity in seafloor sediments and contribute to a variety of essential ecosystem functions including acceleration of organic matter decomposition, bioturbation, and filtration (Covich et al. 1999; Chen 2021). They are an important food source for many species and play a vital role in nutrient cycling and energy flow (Covich et al. 2004; Belley and Snelgrove 2016; Snelgrove et al. 2018; Drejou et al. 2021). Given these attributes they are common biological indicators used in ecological assessments of coastal environments worldwide (Borja et al. 2000; Van Hoey et al. 2010; Ramey et al. 2011).

The distribution and community structure of macrofauna are influenced by several abiotic and biotic processes. Given the intimate association that soft sediment macrofauna have with their sedimentary habitat, differences in sediment grain size, organic content, stability, and porosity all contribute to the well-known, characteristically patchy distributions of benthic communities (Rhoads and Young 1970; Snelgrove and Butman 1994; Snelgrove et al. 2018). Hydrodynamic conditions or water velocity near the seafloor can influence sediment type. Fine sediments (silt; 3.9–63 μm grain size) generally occur in areas of weak flow and allow for greater deposition of organic material, whereas coarser sediments (sand; 63–2000 μm grain size) are present in fast flowing waters with frequent sediment resuspension (Dyer 1986; Ramey and Bodnar 2008). Fine sediments consequently tend to be richer in organic content resulting from both decreased water velocity, as well as their enhanced ability to bind organic carbon (Snelgrove and Butman 1994). Feeding mode of macrofauna is also influenced by hydrodynamics. Generally, deposit feeders have greater abundances in silty/muddy sediments, whereas suspension feeders are associated with coarse sands (Snelgrove and Butman 1994).

Large scale, oceanographic conditions (kms to 100s m) such as temperature, salinity, surface productivity, and circulation are also important factors influencing coastal benthic communities (e.g., Barry and Dayton 1991; Morrisey et al. 1992; Olafasson et al. 1994; Lima and Whether 2012; Alexander et al. 2018). A recent study examining warming based on sea surface temperature (SST) anomalies, in the north Atlantic (i.e., at St. John’s, Newfoundland [NL] Canada) projected an increase of 0.4–2.2 °C over the next 50 years (Han et al. 2015). Spatio-temporal SST gradients are important in influencing wind patterns, storm development, and wave activity through ocean-atmosphere interactions (Bengtsson et al. 2006; Reguero et al. 2019). Attributed to rising SST (C2ES 2020), the North Atlantic Basin has seen an increase in the intensity of tropical storms including hurricanes (Méndez-Tejeda and Hernández-Ayla 2023). Moreover, wave activity in terms of wave height has also increased at high latitudes in recent decades (Reguero et al. 2019). Physical disturbance to the benthic habitat by storms and wave action have the potential to influence seafloor substates (Herkul et al. 2016) through erosion and deposition of sediments. Disturbance from storms can also affect circulation patterns (Ma et al. 2017), salinity, and concentrations of dissolved oxygen (Montagna 2023). Lower salinity and dissolved oxygen caused by storm events can affect the abundance and diversity of benthic macrofauna with recovery times ranging from 3 months to 3 years (Mallin et al. 2002; Montagna 2023). For example, in San Antonio Bay Texas, Montagna (2023) observed a 71% decline in macrofaunal abundance and 54% decline in species/taxon richness after Hurricane Harvey impacted Texas as a category 4 storm.

Changes in environment, especially in terms of sediment type and food resources, can in turn affect the essential ecosystem functions supplied by macrofaunal communities through differences in the biological traits of the different taxa present in the community (Rand et al. 2018; Sutton et al. 2020). Overall, biodiversity (e.g., measured as species richness) has been recognized as an important factor in maintaining ecosystem functioning and the many services associated with the coastal environment (Balvanera et al. 2006; Muntadas et al. 2016). Biodiversity loss can reduce a community’s ability to recycle biologically essential nutrients emphasizing the need to understand the link between biodiversity and ecosystem functioning (Cardinale et al. 2012; Belley and Snelgrove 2016). A functional trait approach, which characterises communities based on the functional roles of species rather than taxonomic identity, can allow for a better understanding of how communities use resources as well as their resilience to environmental change (Sutton et al. 2020). For instance, trait redundancy (e.g., species performing similar functional role) is thought to increase temporal stability by dampening the effect of species loss via function retention (Walker 1992; Biggs et al. 2020). Previous studies have shown that functional richness (i.e., the amount ecological niche space occupied by species in a community) is a better predicator of benthic flux (i.e., changes in oxygen and nutrients) than species richness (Belley and Snelgrove 2016).

Using historical and contemporary benthic macrofaunal composition and abundance data (1998 vs. 2019/2020 respectively) taken 21 years apart, the main objective of the present study was to investigate changes in community and functional structure in Placentia Bay NL and examine environmental drivers of the observed variation. It was hypothesized that historical and contemporary communities in the bay would be different and community patterns would be related to the sedimentary habitat, as it can be influenced by larger-scale oceanographic features. It was expected that functional structure (macrofaunal trait expression) would be related to community structure (taxonomic composition and abundance) such that they would exhibit similar spatio-temporal patterns of variability. It was also hypothesized that there would not be a loss of functional traits/modalities between historical and contemporary communities. Therefore, it was predicted that while species/taxon composition may change, functional roles should remain fulfilled (i.e., no loss of modalities) and observed spatio-temporal patterns based on functional compared to taxonomic composition (i.e., presence/absence) would differ.

## Methods

### Study area

Placentia Bay is the largest embayment on the southeast coast of Newfoundland (47.1030 °N, 54.1859 °W) (Fig. 1). The bay is ∼130 km long and 100 km wide at its southerly directed mouth that is open to the Atlantic Ocean. The average depth in the bay is 125 m (Ma et al. 2012). The inner part of the bay contains three channels divided longitudinally by several islands (Ramey and Snelgrove 2003). Bottom salinity in the bay ranges from 32.2–33.0 (Ramey 2001; Ramey and Snelgrove 2003). Currents are generally influenced by local factors such as wind, water density, and seafloor topography (Bradbury et al. 2000; Ma et al. 2012). In the spring and fall, prevailing winds are from southwest and west, whereas in the winter they are northwest to west (Ma et al. 2012) which can have an affect on surface productivity (van Ruth et al. 2010). Prior to the present study, macrofaunal communities in Placentia Bay were only examined in 1998 when Ramey and Snelgrove (2002) conducted the first comprehensive sampling at ten sites distributed throughout different regions of the bay and on the shelf (hereafter referred to as “historical”: Ramey 2001; Ramey and Snelgrove 2003). Eight of these sites were sampled as part of the present study in 2019 and 2020 (hereafter referred to as “contemporary”) (Fig. 1).

**Fig. 1.**
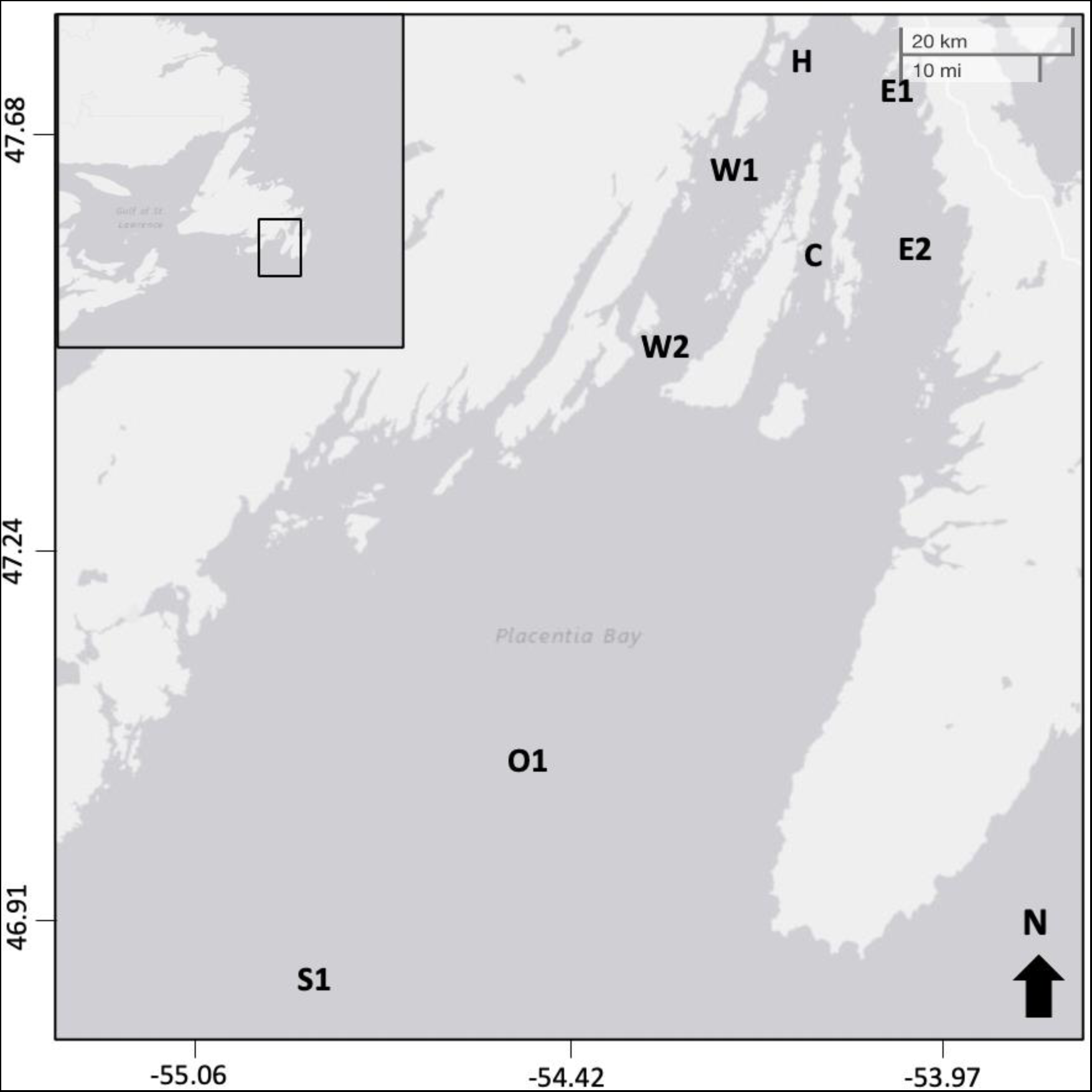
Placentia Bay, Newfoundland showing sampling sites H1 (head of bay), E1/E2 (eastern channel), W1/W2 (western channel), C1 (central channel), O1 (outer bay), and S1(continental shelf), also see Table 1. Inset shows Placentia Bay in relation to Newfoundland.

**Table 1.** Historical and contemporary sampling sites (site/code) indicating latitude and longitude (decimal degrees), mean depth ±SE, and number of samples (no.). Total number samples and area of sea floor sampled: historical: *n*=41, 1.64 m^2^; contemporary: *n*=36, 3.6 m^2^ or 1.8 m^2^/year.

| Site/code | Latitude °N | Longitude °W | Depth m | Historical no. | Contemporary total no. (2019/2020) |
| --- | --- | --- | --- | --- | --- |
| Head/H | 47.7543 | -54.2345 | 63±4.86 | 6 | 6 (3/3) |
| Central/C | 47.5803 | -54.1247 | 202±3.82 | 6 | 6 (3/3) |
| East1/E1 | 47.7475 | -54.0635 | 233±3.36 | 6 | 6 (3/3) |
| East2/E2 | 47.5633 | -54.0428 | 219.5±0.22 | 6 | 3 (0/3) |
| West1/W1 | 47.6448 | -54.2660 | 217±1.49 | 6 | 6 (3/3) |
| West2/W2 | 47.4482 | -54.3298 | 269±17.85 | 6 | 3 (0/3) |
| Outer1/O1 | 47.1800 | -54.3733 | 232±0.67 | 3 | 3 (3/0) |
| Shelf1/S1 | 47.7250 | -54.7967 | 231±1.40 | 2 | 3 (3/0) |

### Sample Collection

Contemporary sample collection was conducted in 2019 and 2020 (i.e., 13–17^th^ September 2019 and 13–17^th^ October 2020) at eight sites in the bay using a 0.1 m^2^ Van Veen grab (Department of Fisheries and Oceans, Canada, NL) (Fig. 1). Three to six independent grab samples were haphazardly collected at each site (total *n*=36), (Table 1). Grab samples were subsampled (50 ml of sediment removed) for determination of sedimentary variables including grain size and total organic matter (top 5 and 1 cm of sediment respectively) (*n*=36). Historical samples used for comparison were collected in 1998 (June and July), from the same eight sites using a random, nested design (Ramey 2001; Ramey and Snelgrove 2003) (Fig. 1, Table 1).

Samples were obtained with a rectangular single-spade box corer containing six subcores. A total of six box cores were collected at each site in the bay, with fewer samples in the outer bay and shelf (total *n*=41; Table 1). For each box core, the four subcores (10 cm × 10 cm wide to a depth of 10 cm; 4x100 cm^2^=400 cm^2^) were used to examine the macrofaunal community and one was used for sediment analyses (Ramey and Snelgrove 2003). Sediment cores were subsampled for total organic carbon (TOC) (i.e., all cores, *n*=38) and grain size (i.e., three random cores per site, *n*=22) (Ramey and Snelgrove 2003). The four macrofaunal subcores were pooled for community composition analysis since they were not considered independent given that they were not completely separated from each other (Ramey and Snelgrove 2003). Any Van Veen and box core samples that were washed out or where the top sediment layer appeared to be disturbed were not retained and were re-sampled.

### Sample processing

All macrofaunal samples were fixed in 4% buffered formalin and then transferred to 70% denatured ethanol with Rose Bengal and sieved over a 500 µm mesh screen prior to sorting macrofauna from the sediments using a stereomicroscope (all years). Macrofauna were counted and identified to the lowest practical taxonomic level, usually species. Taxonomic identifications were completed by L. Treau De Coeli (Université Laval, Quebec) for contemporary samples and P. Ramey-Balci (confirmed by P. Pocklington; Ramey and Snelgrove 2003) for the historical samples.

For grain size analysis, sediments were treated with water/peroxide solution to remove organic material. They were then dried and weighed followed by resuspended and disaggregated prior to sieving to separate the sediment grains into coarse (sand: >63μm), fine (silt: <63–3.9 μm), and clay (<3.9 μm) sized fractions (Wentworth scale), (see Ramey and Snelgrove 2003: historical; Danovaro et al. 2009 and adapted by Bureau Veritas: contemporary). The percent sand, silt, and clay in each sample was calculated using dry weights.

For determination of total organic matter (TOM) of contemporary samples, 50–100 mg of dried sediment was weighed and placed in a muffle furnace for 4 hours at 450°C. The TOM was the difference between dry and calcinated sediments which was normalized and expressed as a percentage (Danovaro 2009). For the historical samples, total organic carbon (TOC) values from Snelgrove and Ramey 2003 were multiplied by 1.724 to determine TOM, as this is a commonly used conversion factor based on the assumption that most soils consist of ∼58% organic carbon (Nelson and Sommers 1996; Pribyl 2010).

### Data Analysis

#### Macrofaunal data

Only infaunal, soft-sediment macrofauna (organisms >500 µm) were included for community analyses following Gallagher and Grassle (1997) and Taghon et al. (2017a). Hard substrate epifaunal encrusting species (e.g., bryozoans, mussels, and hydrozoans) and highly mobile species (e.g., mysids and decapods) not reliably sampled using a Van Veen grab or box corer were also excluded (Taghon et al. 2017a). To take a conservative approach in comparing the historical and contemporary community structure, rare species (species/taxa with <10 individuals across all years and samples) were also removed (Taghon et al. 2017a). The taxonomic status of species was checked against the continuously updated list in the World Register of Marine Species (http://www.marinespecies.org/index.php). To ensure consistent species/taxon identifications between the contemporary and historical datasets, reference specimens (i.e., 48 commonly abundant taxa: P. Ramey-Balci pers. Collection), and identification notes and images contained in Ramey et al. (2001) were compared with contemporary reference specimens. This ensured that species/taxon differences between the two data sets were not a result of discrepancies in identification. On occasion, when identifications could not be confirmed and level of taxonomic identification varied between the two datasets, taxon/species identifications were grouped to a common higher level of classification. The sample size/amount of sediment collected by the different gear types varied for the historical versus the contemporary samples (i.e., historical: box corer, 0.04 m^2^; contemporary: Van Veen grab, 0.1 m^2^) and therefore species densities for the historical data were re-scaled from 0.04 m^2^ to 0.1 m^2^ for data analyses.

#### Functional Traits

A total of eight traits subdivided into 36 modalities were used to describe the functional roles of macrofauna in the bay (Table S1). The functional traits selected describe morphological (i.e., body size), behavioral (i.e., adult movement, bioturbation, living habitat), life history (i.e., reproduction, larval development), and physiological (i.e., feeding mode, tolerance) attributes of macrofauna that directly or indirectly play a role in important ecosystem functions such as nutrient cycling and sediment transport (Martini et al. 2020). Species/taxa for a given modality were scored using the “Fuzzy coding” procedure (Chevenet et al. 1994; Bremner et al. 2006; Oug et al. 2012) based on the following scale, where 0=no affinity (modality not employed), 1=low affinity, 2=high affinity, and 3=exclusive affinity (Oug et al. 2012). For example, when looking at a trait such as feeding mode, species can have more than one modality (e.g., deposit and surface feeding), in this case “1” would be assigned to the less commonly employed modality and “2” to the more commonly employed modality (Oug et al. 2012). If the species/taxon only used one feeding mode (e.g., deposit feeding) a value of 3 was assigned. Modality scores for each trait were assigned based on available databases including the Arctic Trait Base (https://arctictraits.univie.ac.at/), which contained the large majority of the species/taxa in Placentia Bay consistent with its geographic location, as well as The World Register of Marine Species (WoRMS), and Polytrait database (http://polytraits.lifewatchgreece.eu/). This produced a species by trait/modality matrix whereby the modality scores for the species/taxa were multiplied by their abundance in each sample (no. ind. 0.1 m^-2^), and then summed over all species/taxa to obtain a single value for each modality in each sample producing an abundance weighted, functional trait expression matrix for analysis (Bremner et al. 2006; Oug et al. 2012).

#### Multivariate statistics

Variation in macrofaunal community structure and environmental variables between the historical (1998) and contemporary years (2019/2020) were examined by conducting Principal Coordinates Ordination (PCO) analysis in PRIMER (v7 +PERMANOVA) (Anderson, 2001; Clarke and Gorley 2015; Anderson 2017). For the macrofaunal data, the PCO was conducted on a Bray-Curtis dissimilarity matrix generated from species/taxon composition and abundance (at the level of sample; *n* total=77). Prior to analysis, abundances were fourth root transformed to downweigh the relative influence of the more abundant species/taxa, which would otherwise tend to dominate the dissimilarity matrix (Clarke and Gorley 2015). Bray-Curtis dissimilarity shows the proportion of differences in abundance between the samples where 0 indicates samples are similar and 1 indicates samples are completely dissimilar (Legendre and Legendre 2012). Species/taxa with a correlation coefficient of ≥0.7 (Pearson correlation) with the ordination pattern were overlaid as vectors on the PCO plot, where arrows point in the direction of maximal variation in species abundances, and their length is proportional to their maximal rate of change (Ramette 2007). A Permutational Multivariate Analysis of Variance (PERMANOVA; 9999 permutations) tested the null hypothesis that there is no difference in the centroids (equivalent to mean in univariate analysis) in communities between the historical (*n*=41) vs. contemporary years (*n*=36). Similarity percentage (SIMPER, Clarke 1993) was used to identify the species/taxa contributing to 50% of the dissimilarity between years. These analyses were also conducted to examine the composition of species/taxa (i.e., presence/absence) between years whereby PCO was conducted on a Sorensen resemblance matrix. Note that community metrics and multivariate analysis of species/taxon data for 2019 vs. 2020 indicated that they were not significantly different (P>0.5) (see Komendic unpubl. thesis 2023).

Principal Coordinates Ordination analysis (which is equivalent to PCA when based on Euclidean distance) was also used to examine differences among samples based on the sedimentary habitat (i.e., grain size, TOM) and water depth. Draftsman plots (i.e., plots of each variable against each other variable) indicated that environmental variables did not need to be to be transformed prior to analysis, however, all variables were normalized to Z-scores to weigh variables equally (i.e., places them on the same scale). This reduces the degree to which any one variable with a larger mean value sways the result. A PCO analysis was run on this matrix based on Euclidean Distance. A PERMANOVA (9999 permutations) tested the null hypothesis that there is no difference in the sedimentary habitat between the historical vs. contemporary years.

To examine the relationship between the sedimentary environmental variables and macrofaunal data the “Relate” function in PRIMER v7 was used (based on ranks, specifically Spearman rank correlation (ρ) with 999 permutations) (Clarke et al. 2014; Clarke and Gorley 2015). This analysis tested the null hypothesis that there is no correlation between the two matrices (i.e., thus comparing observed patterns in the environmental vs. macrofaunal data). To determine how much of the variation in the macrofaunal data can be explained by the environmental data a distance-based linear model (DistLM) was performed, using the environmental data as a predictor and macrofaunal data/resemblance measures as the response. Predictor variables were added to the model via stepwise procedure and the AICc selection criterion was used to evaluate models.

To compare functional traits between the historical and contemporary years and identify environmental drivers of observed patterns, the same multivariate analyses conducted for the macrofaunal data were repeated for the functional traits with the exception that the PCO was conducted on the Canberra dissimilarity metric generated from the abundance weighted, functional traits expression matrix (instead of Bray-Curtis dissimilarity) to overcome arch effects seen in exploratory analysis via PCO and metric multidimensional scaling (MDS) (Podani and Miklos 2002).

#### Community metrics and univariate statistics

To assess if sampling effort was adequate, species accumulation curves were generated separately for the historical and contemporary samples considering all species/taxa, as well as the major taxonomic groups (i.e., Amphipoda, Bivalvia, Gastropoda, Polychaeta) (PRIMER v7). Pie charts examined the proportion of major taxonomic groups (i.e., percent total abundance) making up the communities in the historical versus contemporary. To compare species composition between time points, the percentage of species/taxa (out of the total species richness for the bay) only sampled in the historical, contemporary, or shared between them were plotted using stacked bar plots. Boxplots compared community metrics including macrofaunal density (no. of ind. 0.1 m^-2^), richness (e.g., number of species/taxa), evenness (Pielou’s Evenness Index, J’), and diversity (Shannon-Wiener, H’ log e) as well as sedimentary variables (i.e., sand, silt, TOC) between years. Student t-tests examined differences in community metrics, abundance of major taxonomic groups, and sediment variables between years. Given the importance that TOM can have in influencing macrofaunal abundance, a linear regression tested if TOM significantly predicted the abundance of polychaetes (given they were the dominant taxon) as well as other major groups (i.e., pooled abundances for amphipods, gastropods, and bivalves) for each timepoint. When the data were normal and had equal variance or could be normalized through transformation (i.e., log or square root) a t-test was directly employed. On occasion when transformations could not normalize the data, a t-test was used when the non-parametric Wilcoxon (Mann-Whitney U) test produced the same result; given that t-tests are robust against non-normality (Rasch et al. 2007). Normality of the distribution of the above datasets was tested using Shapiro–Wilk test and equal variance was examined with Levene’s test of homogeneity. All univariate statistical analyses were performed in JASP (V. 0.16, JASP Team 2022). For functional traits, the proportion of each functional modality (i.e., percent of total expressed traits) as well as the proportions of modalities making up each of the eight traits (such that the values of all modalities for a particular trait add to 100%) were compared for the historical and contemporary samples with the later displayed as circular bar plots (“corrplot” package in R Studio; Wei and Simko 2017).

## Results

### Community structure

The dataset contained 36,909 individuals and 77 species/taxa for analysis. The historical samples included 30,663 individuals (64 species/taxa), whereas the contemporary data contained 6,246 individuals (61 species/taxa). Although more samples were collected in the historical (*n*=41) compared to the contemporary years (*n*=36) the total area of the seafloor sampled was greater for the contemporary (i.e., 1.64 m^2^ vs. 3.6 m^2^ respectively) (Table 1). Overall, the major taxonomic groups including the Amphipoda, Bivalvia, Gastropoda, and Polychaeta made up 90% of the total abundance and >80% of the species/taxa sampled in the bay (historical and contemporary data combined). Species/taxa accumulation curves for the historical and contemporary samples leveled off at ∼36 and ∼29 samples respectively (Fig. 2) and curves for the four major taxonomic groups (with the exception of amphipods) also reached an asymptote (Fig. S1a–d). Principal Coordinates Ordination (PCO) analysis of species/taxon composition and abundance explained 44.2% of the total variability in the data with the historical and contemporary samples generally forming two groups (Fig. 3a). A total of 18 species/taxa including 12 polychaetes and 3 bivalves, contributed ∼50% of the dissimilarity between these time points, with the highest contribution from the polychaete *C. pygodactylata* (6.74%) (Table 2). Note that *C. pygodactylata* was previously identified as *Cossura longocirrata* (see Ramey and Snelgrove 2003). Variation in macrofaunal community structure between the historical vs. contemporary samples indicated they were significantly different (pseudo-F_1, 75_=15.4, PERMANOVA, P=0.0001) (Fig. 3a). Results were consistent even when the dominant species, *C. pygodactylata* was removed from this analysis (PERMANOVA P=0.0001, pseudo-F_1, 75_=13.4). Similarly, PCO analysis of species/taxon composition (i.e., presence/absence; 45.6% of the total variability explained) significantly differed between the historical vs. contemporary samples (PERMANOVA P=0.0001, pseudo-F_1, 75_=13.7) (Fig. 3b).

**Fig. 2.**
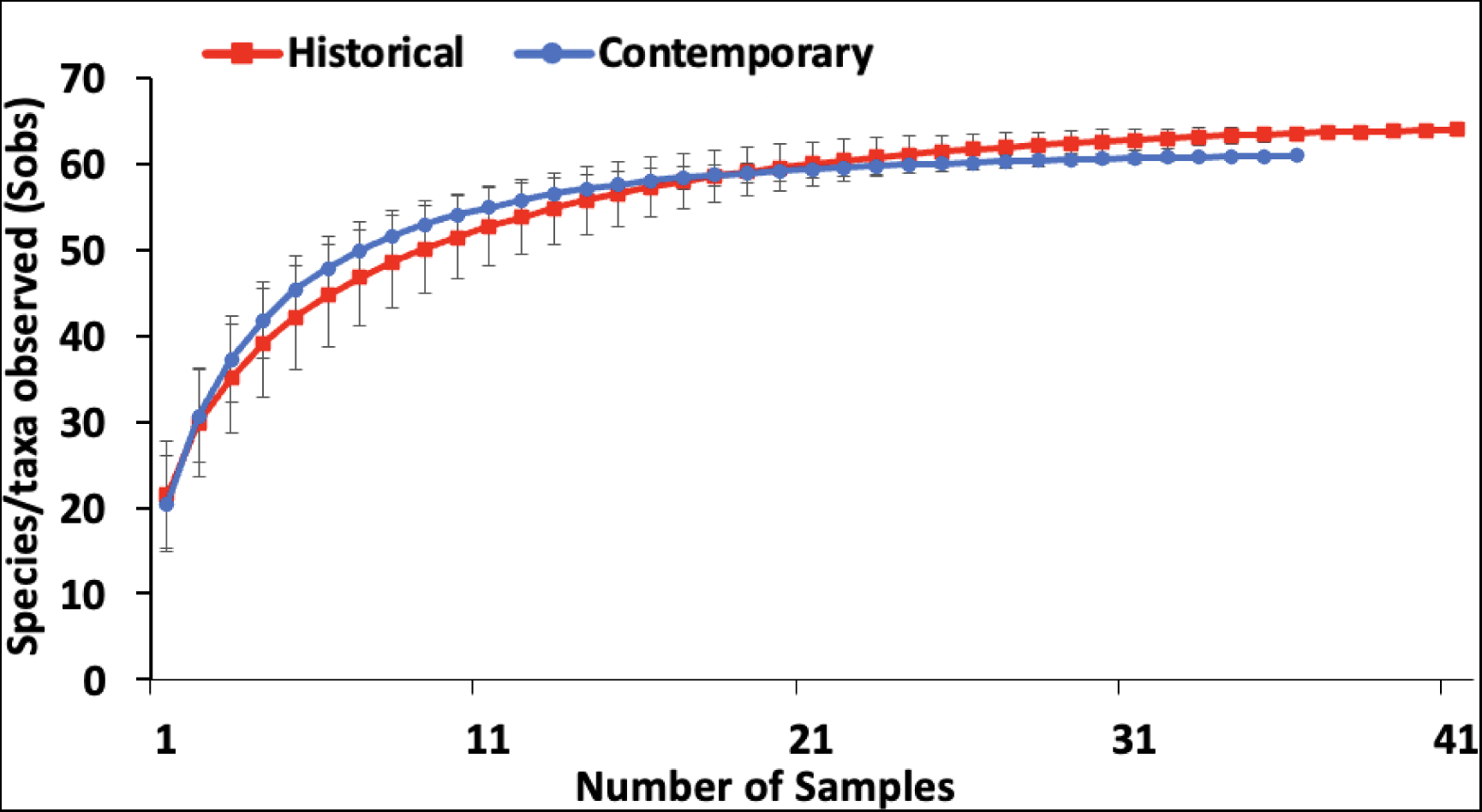
Species accumulation curve ±SD for historical (1998) vs. contemporary (2019–2020) samples.

**Fig. 3.**
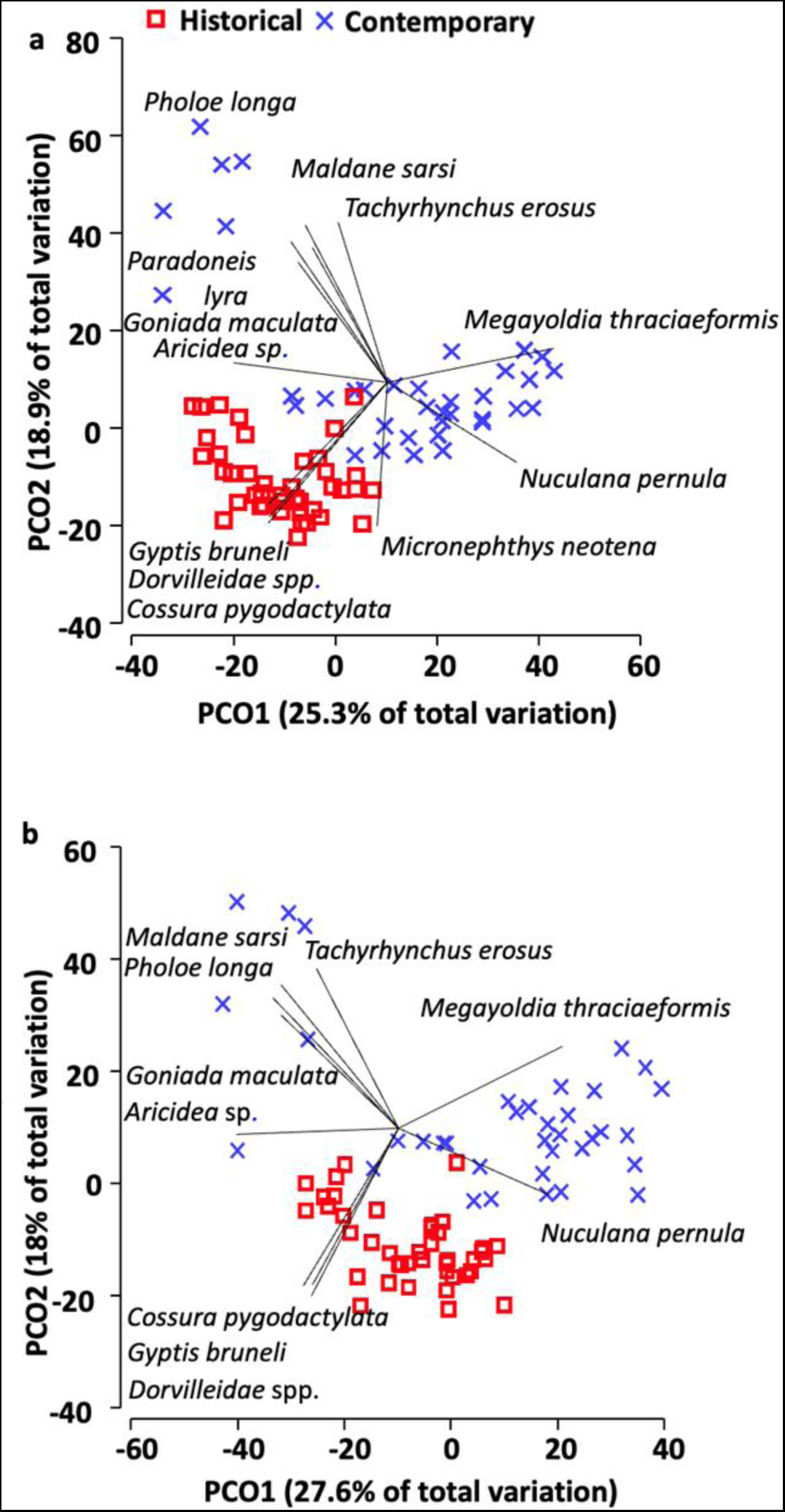
Principal Coordinates Ordination (PCO) of species/taxon (a) composition and abundance based on Bray-Curtis similarity, and (b) presence/absence based on Sorensen resemblance. Differences between the historical versus contemporary communities were statistically significant (pseudo-F_1, 75_=15.4, PERMANOVA P=0.0001 and pseudo-F_1, 75_=13.7, PERMANOVA P=0.0001 respectively). Species vectors=Pearson correlation of ≥ 0.7.

**Table 2.**
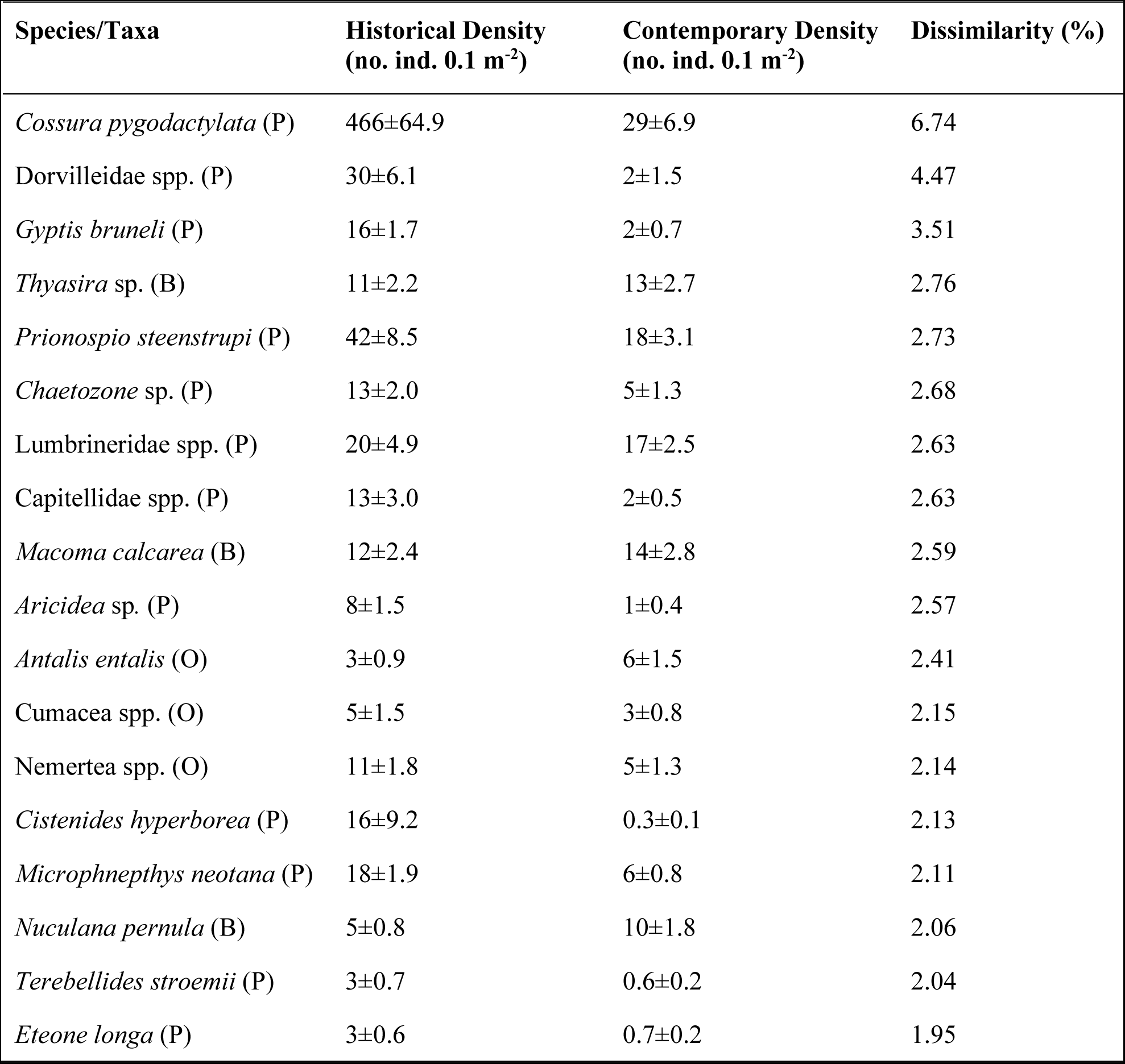
Summary results of SIMPER analysis for species/taxa contributing ∼50% of dissimilarity between historical and contemporary communities including mean density (no ind. 0.1 m**^-2^** ±SE) and their respective contribution to dissimilarity (%). Letters in parenthesis denote P=polychaete, B=Bivalve, O=Other.

Comparison of the major taxonomic groups between the historical and contemporary samples showed the polychaetes made up the greatest proportion of macrofauna collected in both years, however their proportion was much higher in the historical samples (91 vs. 58% respectively) (Fig. 4, also Table S2). In contrast, the proportion of amphipods, bivalves, and gastropods were relatively higher in the contemporary samples (Fig. 4; also Table S2). Moreover, examination of the species making up these major groups found 41 species/taxa (63%) shared between the historical and contemporary years, whereas 14 species/taxa (22%) were only present in the historical samples and 10 species/taxa (15%) were unique to the contemporary (Fig. 5, also Table S3). Mean density of individuals making up the major groups were not significantly different (α>0.05) for the historical vs. contemporary, except for polychaetes which were significantly higher (t_75_=-6.694, P=<0.001) in historical (680 ±80 ind. 0.1m^-2^, *n*=41 vs. 100 ±17 ind. 0.1m^-2^, *n*=36 respectively) (Fig. 6). All community metrics examined, (except for species/taxon richness) differed significantly between years (Fig. 7). Species/taxon evenness (J’) and Shannon diversity (H’) were both significantly lower (t_75_=7.072, p=<0.001 and t_75_=6.38, p=<0.001 respectively) in the historical relative to the contemporary samples (evenness: 0.56 ±0.03, *n*=41 vs. 0.80 ±0.01, *n*=36; diversity: 1.7 ±0.09, *n*= 41 vs. 2.40 ±0.05, *n*=36; respectively) (Fig. 7). Macrofaunal density was significantly higher (t_75_=-7.691, p<0.001) in the historical (748 ±85 no. ind. 0.1 m^-2^, *n*=41) vs. the contemporary samples (174 ±20 no. ind. 0.1 m^-2^, *n*=36) (Fig. 7).

**Fig. 4.**
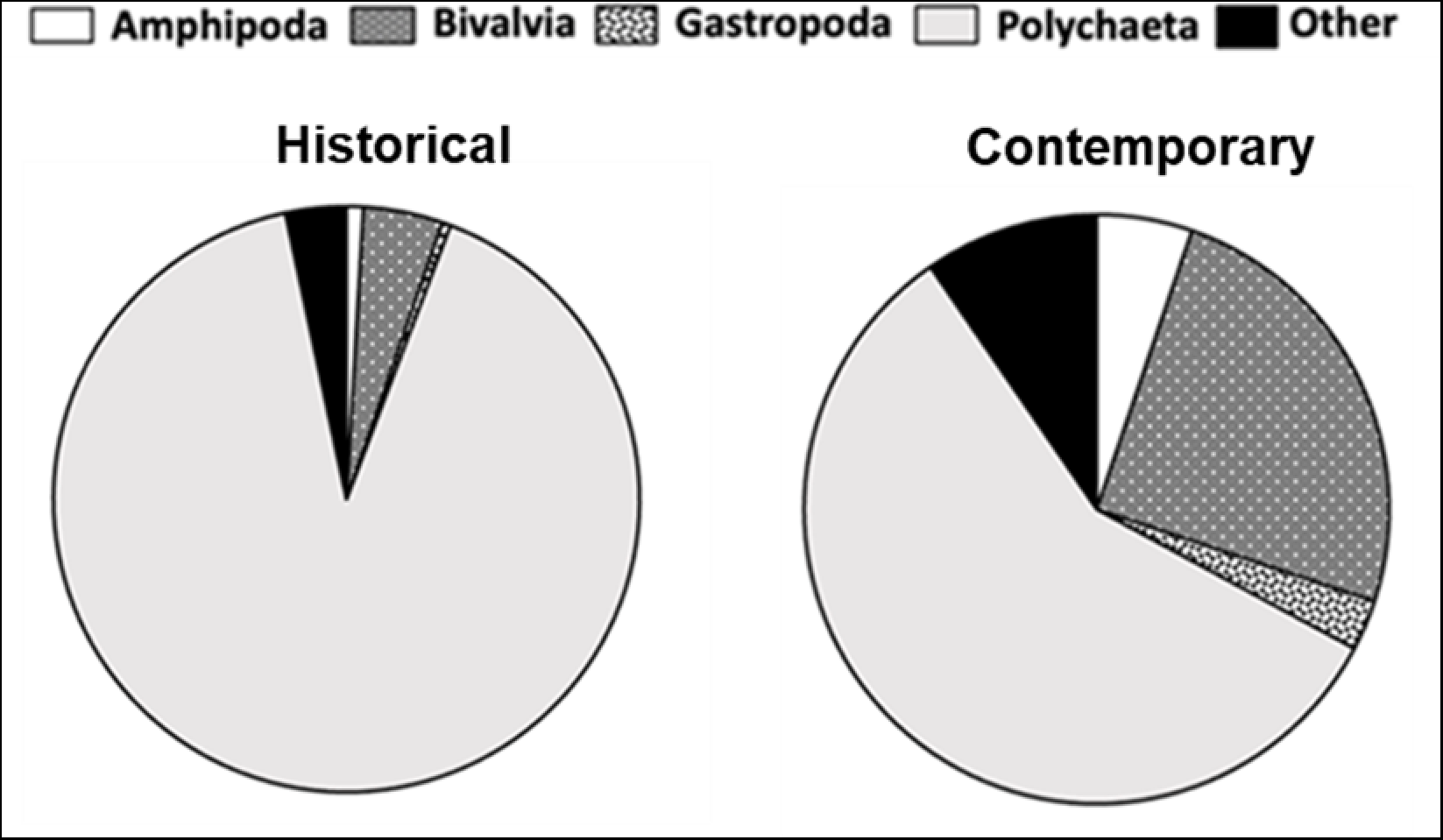
Pie charts showing relative abundance (%) of major taxonomic groups (Amphipoda, Bivalvia, Gastropoda, Polychaeta) in historical versus contemporary communities. Other=remaining taxonomic groups present (e.g., Nemertea and Cumacea).

**Fig. 5.**
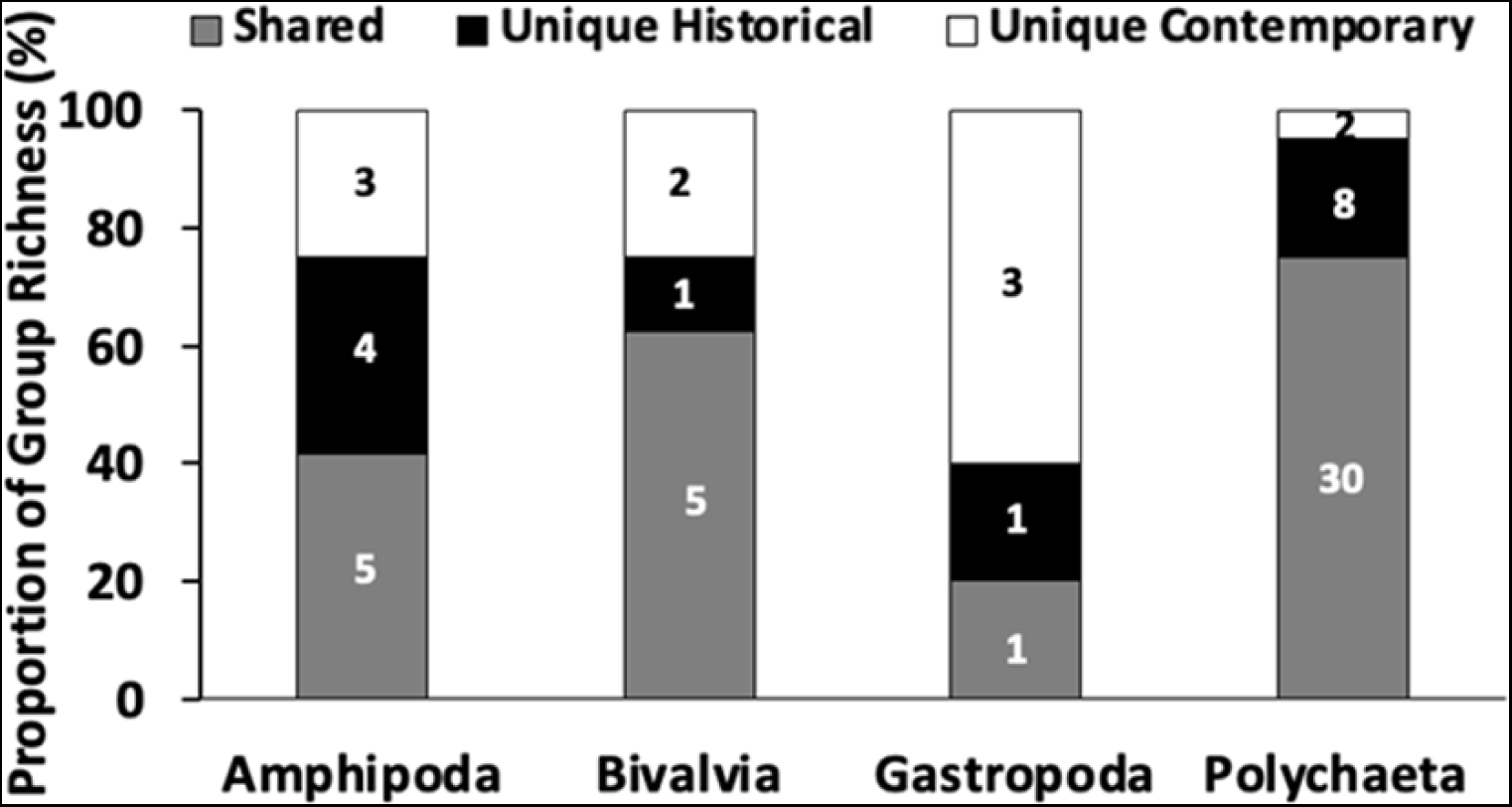
Proportion (%) of total number of species/taxa in samples that were unique to historical (black fill) or contemporary (white) samples or shared (gray) between them for each major taxonomic group (i.e., Amphipoda, Bivalvia, Gastropoda, and Polychaeta). Numbers on bars indicate number of species/taxa shared or unique to each time point.

**Fig. 6.**
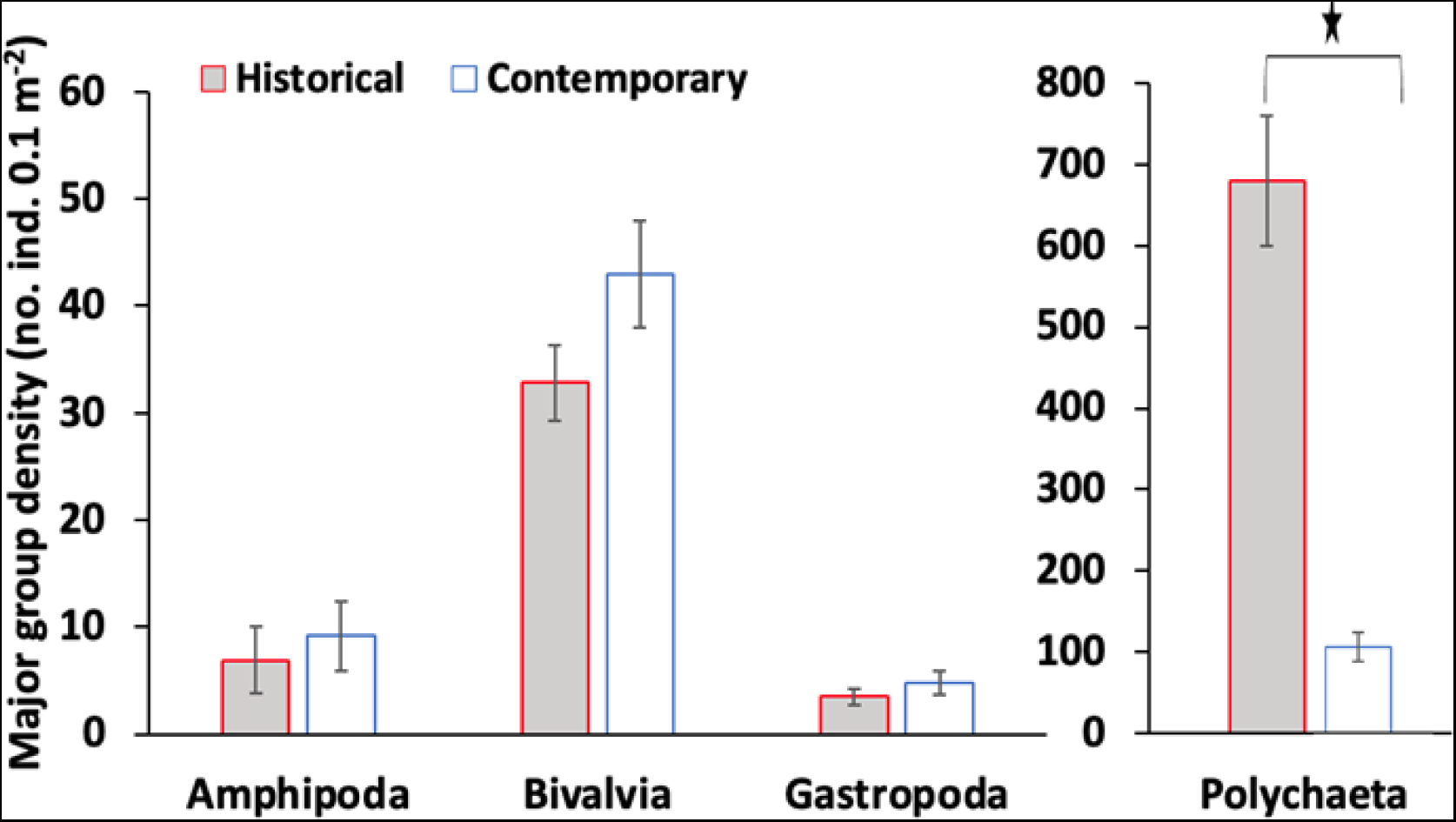
Density (no. of ind. 0.1 m^-2^ ±SE) of major groups in historical (gray) (*n*=41) vs. contemporary (white) (*n*=36) communities. *= significantly different α<0.05.

**Fig. 7.**
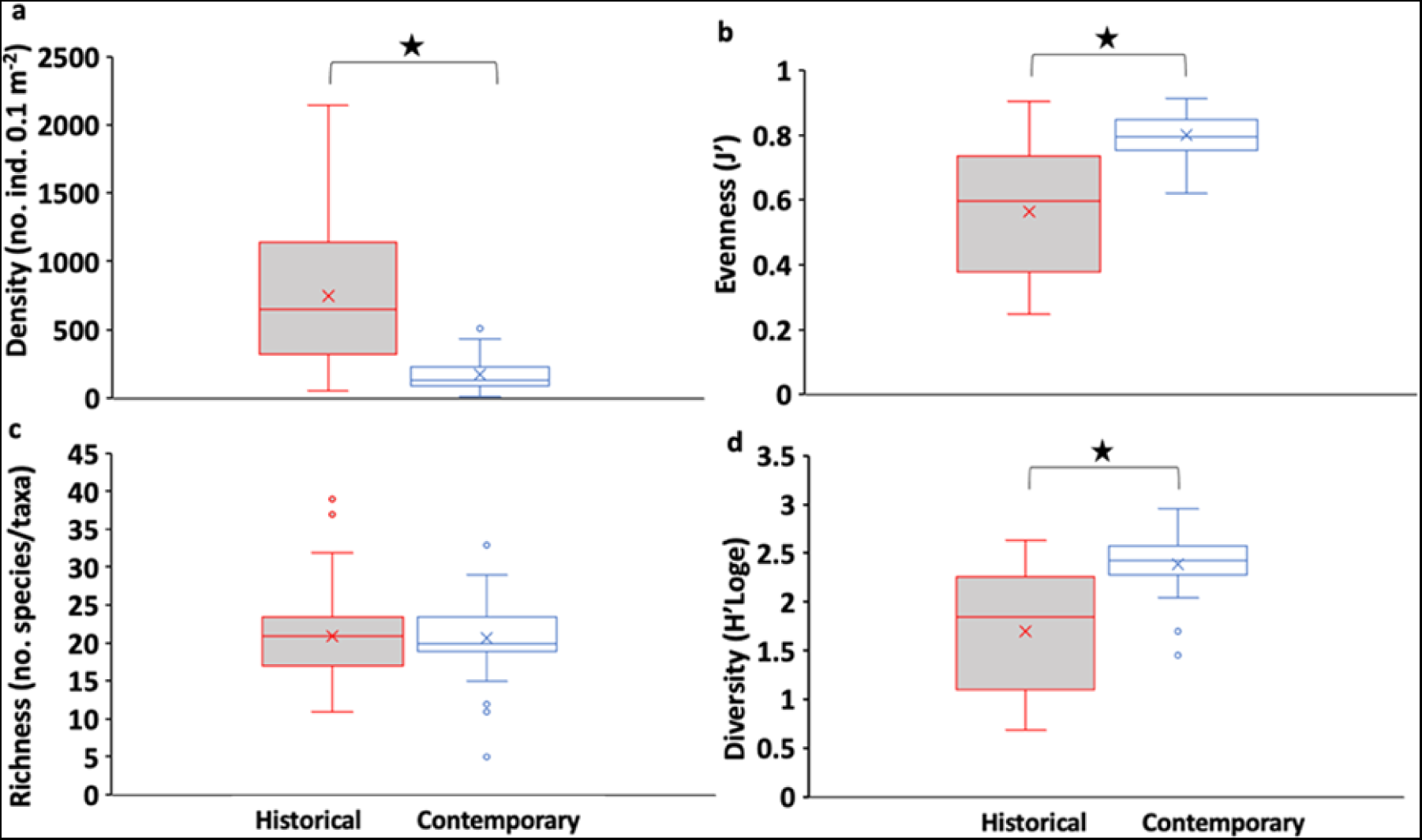
Boxplots comparing (a) density (total number of individuals 0.1m^-2^), (b) evenness (J’), (c) species richness (number of species/taxa 0.1m^-2^), and (d) Shannon diversity (H’log e) for historical (*n*=41) versus contemporary (*n*=36) and communities. Line dividing the box in half represents median and X represents mean. Vertical spread of the box depicts interquartile range encompassing the middle 50% of data. Dots denote outliers. *= significantly different α<0.05.

### Environmental data

Principal Coordinates Ordination (PCO) analysis of the environmental variables explained 78.7% of the total variability in the data. The sedimentary environment significantly differed (PERMANOVA P=0.0001, pseudo-F_1, 55_=60.7) between these two time points (Fig. 8). Examination of sediment grain size showed that the proportion of silt was significantly higher (t_55_=8.6, P=<0.001) in the historical (58 ±2.3%, *n*=22) compared to the contemporary samples (37 ±1.3%, *n*=35), whereas the proportion of sand in the historical samples was significantly lower (t_55_=9.0, P=<0.001) (6 ±1.4%, *n*=22 vs. 27 ±1.5%, *n*=35) (Fig. 9). Total organic matter was also significantly lower (t_72_=-2.8, P=0.006) in the historical (10.4 ±0.76%, *n*=38) compared to contemporary samples (13.5 ± 0.98%, *n*=36) (Fig. 9).

**Fig. 8.**
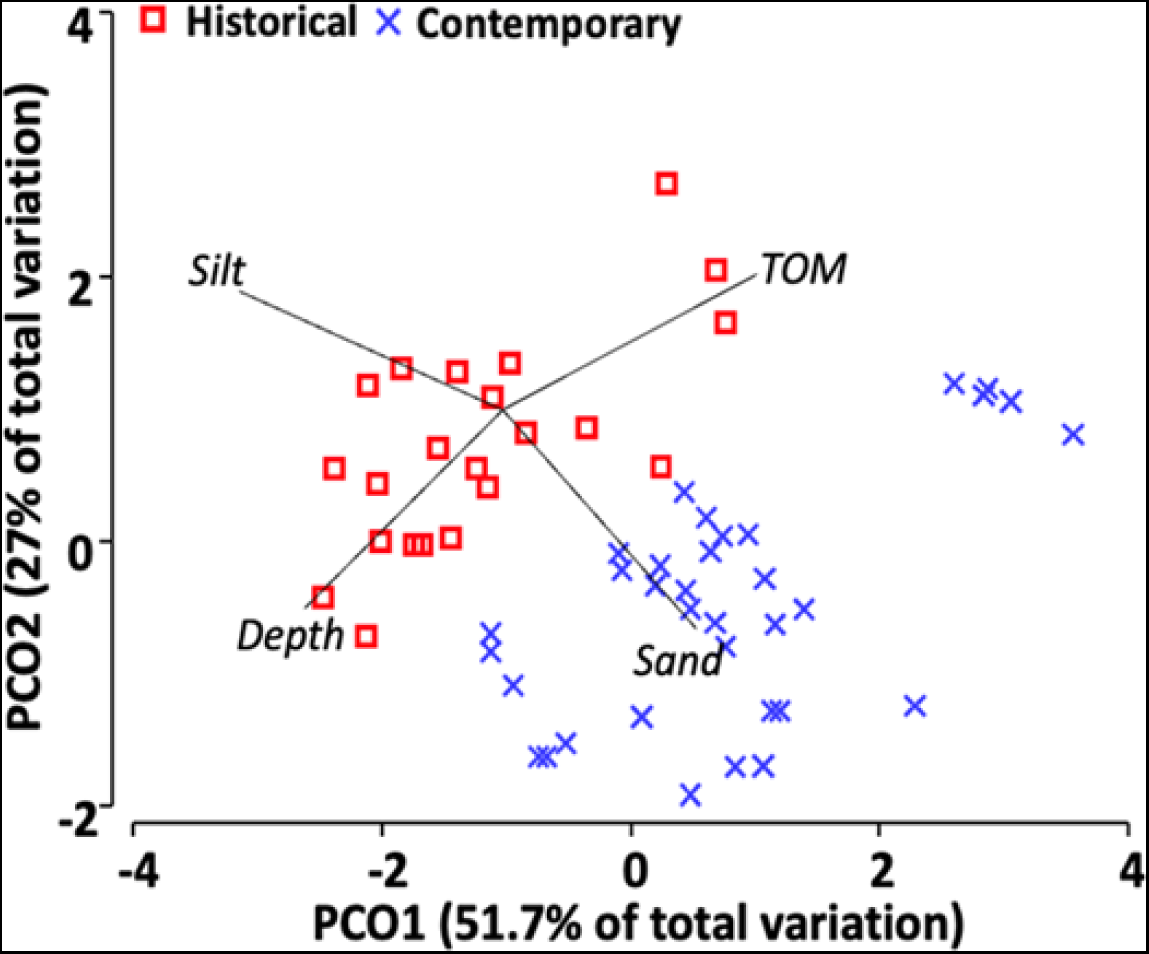
Principal Coordinates Ordination (PCO) of environmental variables (silt, sand, total organic matter, and depth) based on Euclidean distance for historical versus contemporary samples. Differences between the historical and contemporary communities were statistically (PERMANOVA P=0.0001, pseudo-F1, 55=60.7). Abiotic vectors based on Pearson correlation of ≥ 0.7.

**Fig. 9.**
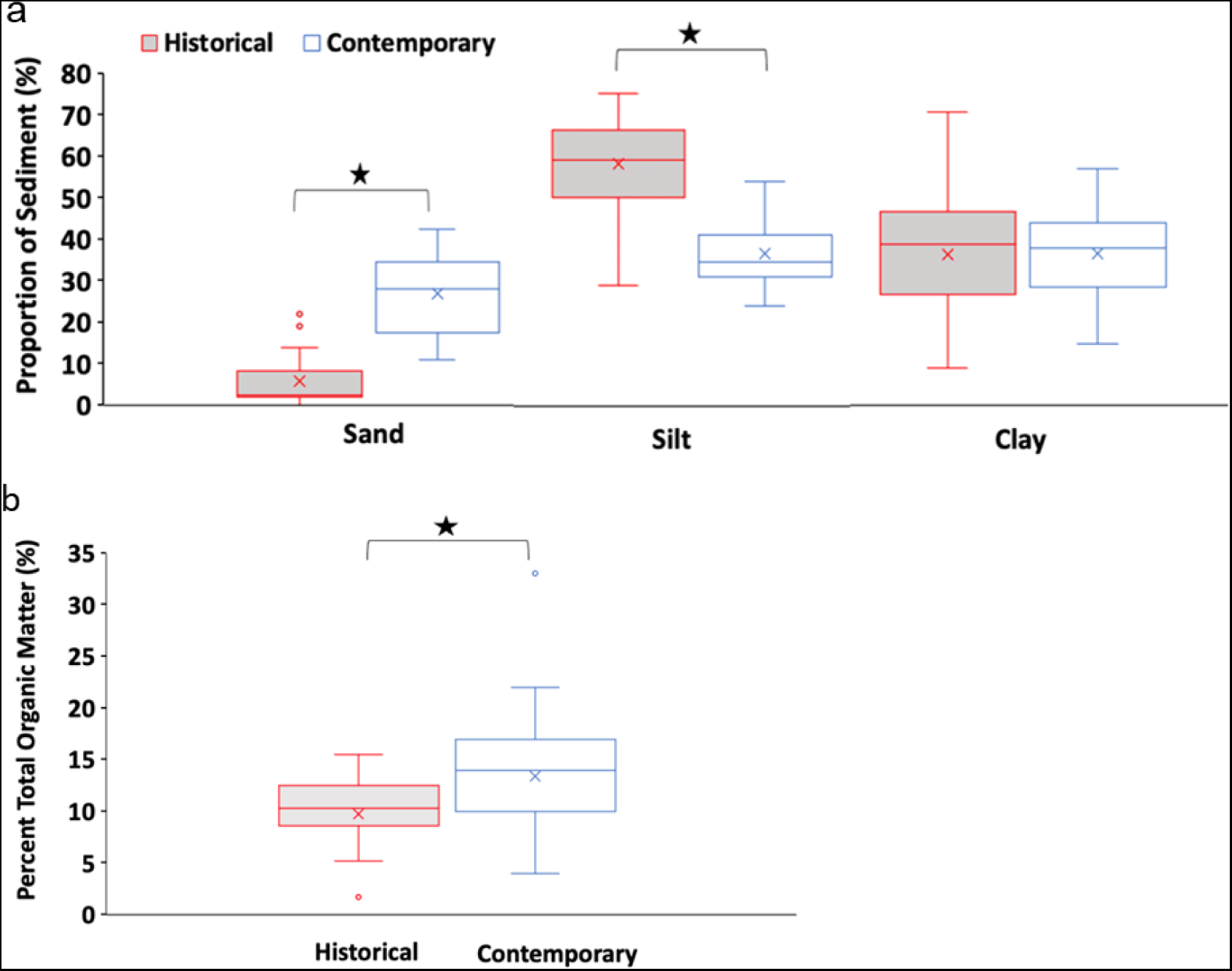
Boxplots comparing (a) sediment grain size (i.e., percent sand, silt, and clay) and (b) total organic matter historical versus contemporary samples. Line dividing the box in half represents median and X represents mean. Vertical spread of the box depicts interquartile range encompassing the middle 50% of data. Dots denote outliers. *=significantly different α<0.05.

### Functional Traits

Principal Coordinates Ordination (PCO) analysis for the expression of functional traits explained 75.2% of the total variability in the data (Fig. S2a). The expressed functional traits significantly differed (PERMANOVA, P=0.0001, pseudo-F_1, 75_=19.5) between the historical and contemporary samples and the spatio-temporal patterns were similar to those observed for species/taxon abundances (RELATE test, ρ=0.554, P=0.001). Ten modalities related to tolerance, body size, feeding, bioturbation, reproduction/larval development, adult movement, and living habitat contributed to ∼50% of dissimilarity between the two time points (Table S4). Examination of modalities making up at least 3% of the total expressed traits for either of the two time points, showed that historical samples contained a relatively higher proportion of highly tolerant (8% vs. 4%, historical vs. contemporary respectively), small body sized (<10 mm, 9% vs. 4%), subsurface deposit feeders (9 vs. 4%), surface deposit bioturbators (8% vs. 5%), and direct developers (9 vs. 5%) (Fig. 10). These modalities, however, were still proportionately most abundant in the contemporary samples, except for small-medium body size and surface deposit feeders (Fig. 10). Overall, examination of the relative proportion of different modalities making up each trait, showed that the reduced expression of dominant modalities in the contemporary samples (i.e., modalities comprising >50% of the total expressed modalities for their respective trait in the historical), allowed for a more even distribution of modalities among the respective traits compared to the historical time point (Fig 10b). In contrast, the composition of functional traits (i.e., presence/absence of modalities) between the two years was similar (PERMANOVA, P=0.250, pseudo-F1, 75=1.6) (Fig. S2b).

**Fig. 10.**
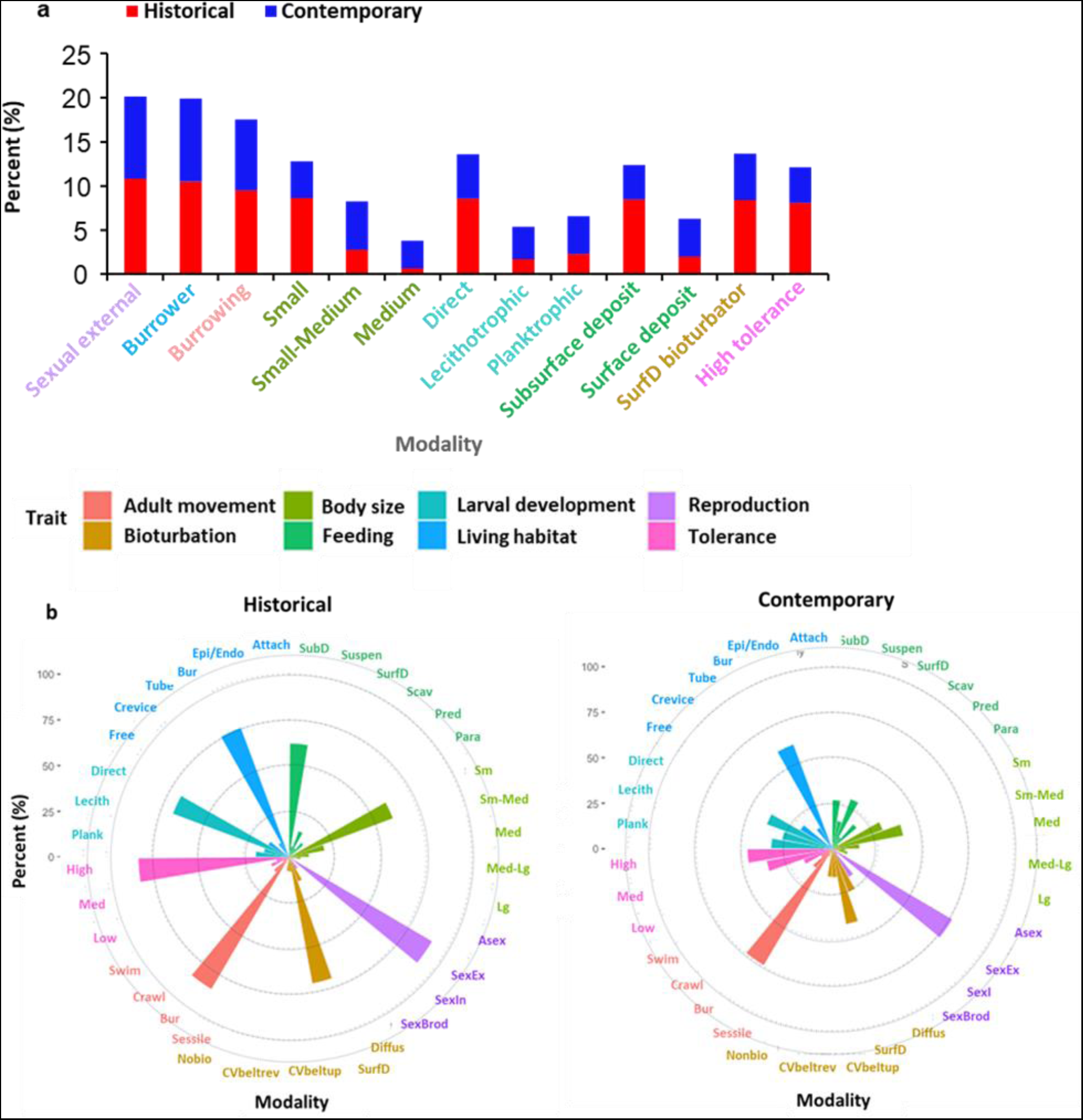
Comparison of functional traits between the historical versus contemporary samples (a) stacked bar plots for modalities making up at least 3% of the total expressed traits, and (b) proportion (%) of modalities making up each of the eight traits. Trait modality definitions defined in Table S1. Note: for (b) modalities for each trait sum to 100%.

### Environmental drivers

Patterns observed in the macrofaunal resemblance matrix (Bray-Curtis dissimilarity generated from species/taxon abundances) significantly matched those found in the environmental resemblance matrix (RELATE test, ρ=0.381, P=0.001). The DistLM indicated that the combination of silt, sand, and TOM were significant in predicting patterns in species/taxon abundance (25% of variance explained) and composition (i.e., presence/absence; 23% of variance explained) (Table 3). Overall, silt explained the largest proportion of the variance (15 and 13% respectively). Patterns observed in the functional trait resemblance matrix (Canberra metric generated from expression of functional traits) were also significantly correlated to those found in the environment resemblance matrix (RELATE test, ρ=0.246, P=0.001). Silt and sand were significant in predicting observed patterns with silt explaining the largest proportion of the variance 23% (total explained 27%) (Table 3). In contrast, patterns based on the functional trait resemblance matrix (Sorensen; generated from presence/absence of modalities) did not significantly match those observed in the environmental resemblance matrix (RELATE test, ρ=0.01, P=0.54) and the environmental variables were not found to be significant predictors of the presence/absence of functional traits. For both historical and contemporary samples, a linear regression indicated that TOM significantly predicted polychaete abundance (R^2^=0.26, F_1,36_=12.9, P=<0.001 and R^2^=0.14, F_1,33_=5.4, P=0.027 respectively) (Fig. S3a) as well as the abundance of other major groups (i.e., pooled abundances for amphipods, gastropods, and bivalves) (R^2^=0.15, F_1,36_=6.3, P=0.017 and R^2^=0.34, F_1,33_=17.1, P=<0.001, respectively) (Fig. S3b).

**Table 3.** Results of Distance Based Linear Model (DistLM) for species/taxon abundance, composition (presence/absence), and expression of functional traits with environmental variables (i.e., sand, silt, total organic matter) as predictors. *=significant predictor

| **Data/**<br>**Resemblance** | **Variable(s)** | **AICc** | **SS**<br>**(trace)** | **Pseudo-**<br>**F** | **P-value** | **Prop.**<br>**R<sup>2</sup>** | **Cumul.**<br>**R<sup>2</sup>** | **Res.**<br>**DF** |
| --- | --- | --- | --- | --- | --- | --- | --- | --- |
| Abundance:<br>Bray-Curtis | silt | 417.4 | 13627 | 9.36 | *0.001 | 0.15 | 0.15 | 55 |
|  | sand | 415.0 | 6156 | 4.50 | *0.001 | 0.07 | 0.21 | 54 |
|  | TOM | 414.7 | 3371 | 2.53 | *0.016 | 0.04 | 0.25 | 53 |
| Composition:<br>Sorensen | silt | 410.2 | 10550 | 8.24.5 | *0.001 | 0.13 | 0.13 | 55 |
|  | sand | 407.9 | 5389.5 | 2.3 | *0.002 | 0.07 | 0.20 | 54 |
|  | TOC | 407.8 | 2724.2 |  | *0.043 | 0.03 | 0.23 | 53 |
| Expression of<br>functional traits:<br>Canberra | silt | 183.3 | 394 | 16.4 | *0.001 | 0.23 | 0.23 | 55 |
|  | sand | 182.6 | 65 | 2.7 | *0.042 | 0.04 | 0.27 | 54 |

## Discussion

Macrofaunal community and functional structure, and sedimentary habitats in Placentia Bay have undergone significant changes since 1998. Overall, 14 species/taxa were only observed in the historical community, whereas 11 species/taxa were unique to the contemporary. While 63% of species/taxa making up the major taxonomic groups were shared between these two time points, this contrasts with 86% shared between the two contemporary years (2019 vs. 2020) which showed similar spatio-temporal patterns in community structure and community metrics (at α=0.5) (see Komendic, unpub. thesis 2023). The contemporary community exhibited a lower density of macrofauna but had similar richness, resulting in higher evenness and diversity compared to the historical one. Community and functional structure in the bay were related to corresponding changes in the sedimentary habitat, known to be important in structuring benthic communities. Most notably there have been changes in sediment grain size and total organic matter content such that in contemporary years grain size is coarser and organic matter is higher relative to 1998. While spatio-temporal patterns of variability based on functional expression of macrofaunal traits reflected those in taxonomic composition and abundance, functional composition (i.e., presence/absence of modalities/traits) did not vary between the historical and contemporary communities, despite changes in taxonomic composition. Similarly, Sutton et al. (2020) found that epibenthic communities in two Alaskan Arctic shelfs were more similar in terms of their functional and taxonomic diversity than in functional and taxonomic composition. Such findings (including those of the present study) lend some support to the idea that redundant functional traits (implying ecological functioning) may be able to be maintained under changing species composition as taxa being eliminated are replaced by others with similar traits (Frid and Caswell 2015; McLean et al. 2019). However, changes in the relative expression of functional traits are also important to consider as they can influence which resources are used and how effectively different services are able to be maintained (Cadotte et al. 2011; Sutton et a. 2020).

Although temporal samples taken at specific time points such as decades apart have inherent limitations that make short-term vs. longer-term community variability difficult to interpret (Renaud et al. 2007; Herder et al. 2021) they are necessary in understanding and predicting current and future community changes (e.g., Van Geest et al. 2015; Taghon et al. 2017a; Harris et al. 2023; Ramon et al. 2023). Therefore, it is pertinent to consider differences in sampling methods and protocols when making comparisons and interpreting results (Taghon et al. 2017a). In our case, yearly sampling effort and sieve size were consistent among years. Furthermore, taxonomic discrepancies were clarified between the two datasets using existing reference collections. While the grab used to collect the contemporary samples was larger than the box core used for the historical sampling, species/taxon richness was similar and density of macrofauna was lower in the contemporary samples (even before the scaling of historical samples to facilitate comparisons); which is the opposite trend than one might expect based on sample size alone. Species densities determined using a Van Veen grab versus box core samples in fine sands have been shown to be comparable, except for deeper living species when the Van Veen does not penetrate past the top 5 cm of sediment (Beukema 1976). In our study, the average penetration depth of the Van Veen grab was 12.6 ±4.1 cm.

Species accumulation curves generated separately for both time points showed that the overall community as well as the major taxonomic groups were adequately sampled (apart from amphipods). Thus, an increase in species richness would not be expected with additional sampling. We did however have a discrepancy with respect to sampling season which can result in marked changes in species abundances, especially in temperate regions (Van Geest et al. 2015; Taghon et al. 2017a); although other studies have not found changes in macrofaunal abundances with season (e.g., North Atlantic: Valderhaug and Gray 1984; Weissberger et al. 2008). While season in the present study may have contributed to some of the unexplained variability between years, as relatively lower densities may be expected in the fall (sampled in contemporary, Sep/Oct) following the summer reproductive period (historical samples collected in the summer, Jun/Jul) (Reiss and Kröncke 2005), sediment grain size and TOM were significant in predicting in predicting patterns in macrofaunal species/taxon composition and abundance. Moreover, lower abundance in the fall 2019/2020 was taxon specific being only observed for the polychaetes and considerable changes in community composition (i.e., presence/absence of species/taxa) were also observed (even after the removal of rare species); which would be less likely to vary substantially with season.

More specifically, in terms of changes in the abundance of polychaetes, *Cossura pygodactylata,* Dorvilleidae spp., and *Gyptis bruneli* in particular, showed large reductions (>50%) in their mean densities in the contemporary community relative to the historical one. *Cossura pygodactylata* (Family Cossuridae) was the most abundant species present at both time points and this family has been observed to vary seasonally. Weissberger et al. (2008) examining macrofaunal communities in the Gulf of Maine found the density of cossurid polychaetes was lowest in the fall but despite this it remained a dominant taxon. Similarly in the present study, even with the lower density of *C. pygodactylata* in the contemporary community (sampled in the fall), it remained the most dominant species in the bay and community structure between the historical vs. contemporary time points remined statistically different even when it was removed from the multivariate analyses. In contrast, bivalves, amphipods, and gastropods showed the opposite trend, whereby their relative proportions and abundances were generally higher (but not significantly different) in contemporary relative to the historical community. Bivalves were also responsible for some of the observed dissimilarity between the two time points.

The sedimentary habitat in Placentia Bay has also undergone significant changes in sediment type. This contrasts with studies examining changes in sediment type overtime that have indicated that grain size tends to remain relatively stable (Taghon et al. 2017a; Herder et al. 2021). For example, Taghon et al. (2017a) found that sediment properties in Barnegat Bay, New Jersey, showed few changes in 45 years such that the silt/clay content was similar in 1965–69, 2000–06, and 2012–14. Likewise, a previous long-term study in Frobisher Bay, Nunavut, Canada, found that grain size did not significantly change over a 50-year period (i.e., between 1967–1976 and 2016) (Herder et al. 2021). In Placentia Bay, contemporary sediments were ∼4.5x coarser as compared to the historical time point. As physical disturbance created by storms can modify seafloor substates (Herkul et al. 2016) it is plausible that changes in sediment grain size in the bay may be related to storm activity. Disturbance by waves and currents during storm events can cause sediment transport (i.e., deposition), as well as removal (i.e., erosion) in coastal regions (Manchia et al. 2023). While Taghon et al. (2017b) found no change in sediment median particle size after hurricane Sandy, it is possible that sediments in Barnegat Bay were not affected because the bay is protected by two barrier islands that shelter it from the Atlantic Ocean waters. In contrast, Placentia Bay has a large mouth (100 km wide) and no prominent sill, exposing it to the adjacent shelf environment allowing significant exchange of water with the Atlantic Ocean (Maclsaac et al. 2023). Additional sampling in the bay is needed to determine whether the bay has undergone sustained changes in sediment grain size, however, sediments have become consistently coarser at nearly all sites from 2019 to 2021 (Komendić unpubl. thesis 2023). Placement of current meters at some of the established study sites would aid in determining whether bottom flows during storm events is a plausible mechanism for the observed changes in sediment type.

Sediment grain size (i.e., silt and sand) was an important predictor of community and functional structure in the bay (except functional composition) with silt explaining the largest proportion of the observed variation. Overall, fine sediments in the historical community were associated with highly tolerant small bodied subsurface deposit feeders such as *C. pygodactylata*, and Dorvilleidae spp. which contributed to ∼11% of the dissimilarity between the time points. In comparison coarser sediments in the contemporary community contained a relatively higher proportion of small-medium (or medium sized), surface deposit feeders (or suspension feeders) having pelagic planktotrophic or lecithotrophic larvae of which three bivalves (i.e., *Nuculana pernula*, *Macoma calcarea* and *Thyasira* sp.) contributed ∼7% to dissimilarity. Similarly, in the Wadden Sea Gusmao et al. (2022) examining functional traits along a sediment gradient found that coarser sediments were dominated by surface modifiers, suspension feeding organisms, with larger bodies, whereas fine sediments associated with higher levels of organic matter were dominated by deposit feeders with small bodies. Kun et al. (2019) also found functional traits to vary with sediment type in Arctic Bering Sea where silty-sandy sediments contained tube-dweller/burrower, deposit feeding modalities, compared sandy sediments predominantly containing motile taxa.

In addition to changes in sediment grain size, there has also been an increase in sedimentary TOM from the historical to contemporary times (i.e., 10.4 ±0.52% vs. 13.5% ±0.98%, respectively). Though TOM is generally positively related to density (Davies and Payne 1984; Nestlerode et al. 2020), organic loading can decrease macrofaunal abundance (Pearson and Rosenberg 1978; Snelgrove and Butman 1994). In the present study, TOM was found to significantly predict the density of macrofauna at both time points. In the historical community Ramey and Snelgrove (2003) attributed low macrofaunal densities to sulphide production in the sediments as well as poor food quality driven primarily by low densities at three sites which had the highest percentage of TOC. Similarly, darkened sediments (i.e., suggesting hydrogen sulphide accumulation) were seen during contemporary sample collection in both 2019 and 2020. Studies examining the effect of sediment organic carbon content on benthic communities in the Northern hemisphere noted a range from 10–35 mg g^-1^ (which is equal to, TOM: 1.7– 6.0%; TOC: 1.0–3.5%) as the general threshold where benthic communities would be expected to be reduced (Hyland et al. 2005; Walker et al. 2022).

While TOM in Placentia Bay sediments was also important in predicting species/taxon composition and abundance patterns in the bay it explained much less of the variability, and it was not an important predictor of functional structure. Interestingly, levels of TOM were significantly elevated in the coarser contemporary sediments as compared to the finer sediments characterizing the historical time point (i.e., 10.4 ±0.52% vs. 13.5% ±0.98%, respectively). As TOM is commonly associated with finer sediment grain sizes (Snelgrove and Butman 1994), this decoupling may also be suggestive of possible effects of disturbance (e.g., storms and organic enrichment). Freshwater input during storm events, and/or from aquaculture in the bay, may be a source of organic enrichment (Haya et al. 2001; Bêche et al. 2006; Fuch et al. 2020). Supply of organic matter can vary year-to-year, influenced by both freshwater inputs and climate/weather (Diaz and Rosenberg 1995). Placentia Bay experiences heavy rainfall during storm surges (e.g., ∼200 mm hurricane Teddy, September 22, 2020), and has freshwater inputs at the head of the bay including the Come by Chance River (mean discharge∼1.95m^3^s^-1^) (Maclsaac et al. 2023) and Swift Current (Ramey et al. 2003). Moreover, sedimentary TOM has been shown to be highest at the head of the bay compared to other areas (Ramey et al. 2003; Komendić unpubl. thesis 2023). Changes in SST (Han et al. 2015) may also play a role as it can influence timing and amount of food supply (e.g., primary production) to the benthos. Overall, the reduced abundance of highly tolerate, deposit feeding polychaetes in relatively coarser sediments characterizing the contemporary years, appears to support a more even distribution of functional traits related to tolerance, body size, feeding, bioturbation, and larval development which are important in resistance to disturbance, resources use, and energy flow (e.g., Rand et a. 2018; Sutton et al. 2020). Relatively high functional evenness has been suggested to support resilience through its relatively greater potential for maintenance of ecosystem function (Sutton et al. 2020).

## Conclusion

Communities and habitats in Placentia Bay have undergone significant changes since 1998. Overall, the initial hypotheses were consistent with the results. Community and functional structure in the bay significantly differed between historical and contemporary times and were influenced by the sedimentary habitat. Moreover, trait modalities were maintained between these time points (i.e., no loss of modalities), however modalities were more evenly distributed in terms of the proportion of expressed traits in the contemporary community. Our findings suggest that communities have exhibited temporal stability or resilience with respect to functional traits despite the current levels of TOM; possibly through combined affects of high sediment TOC and increased grain size influencing the abundance of highly tolerate deposit feeding polychaetes. Future monitoring in the bay should prioritize sampling to better quantify the extent of year-to-year seasonal variation of macrofauna in relation to the sedimentary habitat (i.e., sediment grain size, TOC, and C/N ratios) at critical time points including the summer and fall months. A long-term sampling station(s) that could be more frequently sampled would be useful in capturing shorter/longer-term changes/impacts associated with disturbances that may result from nearby aquaculture, temperature change, and sporadic events such as large inputs of rainfall, storm surges and hurricanes.

## Acknowledgments

We would like to thank the Benthic Ecology team (DFO-NL) and colleagues for collection of the field samples in 2019–2022 (K. Bussey, R. Deering, R. Dove, L. Freeman, V. Hayes, P. Hawkins, E. Herder, C. Malayny, B. Piercey, and A. Robar), as well as Burin Tradition’s captain W. Pitcher and L. Fudge (vessel crew). L. Treau-De Coli identified the macrofauna. Dr. G. Davoren provided helpful comments on earlier drafts of the manuscript. Research was supported by the Placentia Bay Coastal Environmental Baseline Program (DFO-NL, Canada) and associated colleagues K. Dalley, L. Freeman, J. Higdon, and K. Tucker.

