## Supplementary Tables & Figures for "What has changed in 20 years? Structure and function of soft-sediment macrofauna in a subarctic embayment, Newfoundland (Canada)"

**Supplementary Material**

**Table S1.** Functional traits and modalities with corresponding abbreviations and associated ecosystem function. Definitions based on those provided by the Arctic Trait Base (Degen and Faulwetter 2019).

| Trait | Modality | Definition | Abbreviation | Function |
|---|---|---|---|---|
| Adult Movement | Sessile | No movement | Sessile | Foraging ability, predatory avoidance, dispersal abilities |
| | Burrower | Burrows in sediments | Bur | |
| | Crawler | Move along sediment via legs or other appendages | Crawl | |
| | Swimmer | Moves above the sediment | Swim | |
| Body Size | Small | <10 mm | Sm | Energetic demand, ability to resistance predation |
| | Small-medium | 10—50 mm | Sm-Med | |
| | Medium | 50—100 mm | Med | |
| | Medium-large | 100—300 mm | Med-Lg | |
| | Large | >300 mm | Lg | |
| Larval Development | Pelagic/Planktrophic | Larvae grow in the water column | Plank | Fecundity, development, and dispersion insights |
| | Pelagic/lecithotrophic | Larvae with yolk sac, pelagic period is short | Lecith | |
| | Benthic/direct | Larvae have benthic development | Direct | |
| Reproduction | Asexual | Budding & fission | Asex | Dispersal, continuous reproduction could allow for resilience, production |
| | Sexual: external | External fertilization (e.g., eggs & sperm released into water) | SexEx | |
| | Sexual: internal | Internal fertilization | SexIn | |
| | Sexual: brooding | Internal or external fertilization but eggs are brooded | SexBrod | |
| Bioturbation | Diffusive mixing | Random mixing of particles | Diffus | Impacts on biogeochemical cycles, food acquisition, resistance from disturbance |
| | Surface deposit | Deposition of particles at sediment surface (e.g., from defecation) | SurfD | |
| | Conveyer belt (upward) | Movement of sediment/particles from within sediment to the surface | CVbeltup | |
| | Conveyer belt (reverse) | Movement of sediment/particles from surface to deep within sediments | CVbeltrev | |
| | None | No bioturbation | Nobio | |
| Feeding Mode | Subsurface deposit | Feeds within the sediment | SubD | Method of resource acquisition, living position, growth requirements |
| | Suspension | Feeds on particles suspended in the water | Suspen | |

| | | | | |
|---|---|---|---|---|
| | Surface deposit | Feeds on material from sediment surface | SurfD | |
| | Scavenger | Feeds on a variety of particles | Scav | |
| | Predator | Feeds on other organisms | Pred | |
| | Parasite | Uses host to obtain food | Para | |
| Living Habitat | Free living | Freely move within/on sediments | Free | Position in sediment, food acquisition, preferred environmental conditions |
| | Crevice dwelling | Tend to live in spaces between rocks | Crevice | |
| | Tube dwelling | Create tubes | Tube | |
| | Burrowing | Burrow within sediments | Bur | |
| | Epi/endo/phytic | Live on/in other organisms | Epi/Endo | |
| | Attached | Live attached to substrate | Attach | |
| Tolerance | Low | Species sensitive to changes in environment (e.g., organic enrichment, pollution, temperature, salinity changes) | Low | Species/taxa tolerance to disturbance (e.g., temperature, salinity, organic enrichment). Low indicating very sensitive and high indicated tolerant to conditions |
| | Medium | Species indifferent | Med | |
| | High | Species tolerant | High | |

**Table S2.** Density (no. ind. 0.1m$^{-2}$ ±SE) of species/taxa for major taxonomic groups (i.e., Amphipoda, Bivalvia, Gastropoda, Polychaeta, other) in historical versus contemporary samples.

| Major Group | Species/Taxa | Historical Density (no. ind.m$^{-2}$ ±SE) | Contemporary Density (no. ind.m$^{-2}$ ±SE) |
|---|---|---|---|
| **Amphipoda** | *Aceroides (Aceroides) latipes* | 0.12±0.09 | 0.30±0.14 |
| | *Bathymedon* sp. | 1.40±0.46 | 0.00±0.00 |
| | *Byblis gaimardii* | 0.67±0.40 | 0.09±0.07 |
| | *Hippomedon* sp. | 1.95±1.04 | 0.09±0.07 |
| | *Megamoera dentata* | 0.00±0.00 | 0.27±0.19 |
| | *Melita* sp. | 0.43±0.30 | 0.00±0.00 |
| | *Monocludes* sp. | 0.12±0.085 | 0.51±0.31 |
| | *Orchomenella minuta* | 0.00±0.00 | 0.75±0.23 |
| | *Paratryphosites abyssi* | 0.00±0.00 | 5.59±3.19 |
| | *Pontoporeia femorata* | 1.16±0.50 | 1.56±0.38 |
| | *Protomedeia* sp. | 0.73±0.73 | 0.00±0.00 |
| | *Quasimelita formosa* | 0.31±0.22 | 0.00±0.00 |
| **Bivalvia** | *Axinopsida orbiculata* | 0.00±0.00 | 1.97±1.18 |
| | *Ennucula* sp. | 1.59±0.44 | 0.99±0.24 |
| | *Macoma calcarea* | 12.13±2.37 | 14.11±2.78 |
| | *Megayoldia thraciaeformis* | 0.00±0.00 | 1.63±0.34 |
| | *Nuculana pernula* | 5.18±0.75 | 9.54±1.81 |
| | *Thyasira* sp. | 11.10±2.15 | 13.25 ± 2.71 |
| | *Yoldia hyperborea* | 0.37±0.16 | 1.50±0.31 |
| | *Yoldia* sp. | 2.44±0.60 | 0.00±0.00 |
| **Gastropoda** | *Curtitoma incisula* | 0.00±0.00 | 1.08±0.30 |
| | *Propebla rugulata* | 0.00±0.00 | 1.56±0.38 |
| | *Retusa obtusa* | 2.38±0.60 | 0.63±0.40 |
| | *Tachyrhynchus erosus* | 0.00±0.00 | 1.23±0.46 |

| | | | |
|---|---|---|---|
| | *Turridae* sp. | 1.10±0.34 | 0.00±0.00 |
| **Polychaeta** | Ampharetidae sp. A | 0.92±0.63 | 0.00±0.00 |
| | *Ampharete finmarchia* | 0.24±0.15 | 0.00±0.00 |
| | *Apistocranchus typicus* | 0.79±0.32 | 0.63±0.37 |
| | *Arcteobia anticostiensis* | 0.06±0.06 | 0.54±0.18 |
| | *Aricidae sp.* | 7.80±1.53 | 1.32±0.39 |
| | *Artacama proboscidae* | 1.77±0.84 | 0.00±0.00 |
| | *Bradabyssa villosa* | 1.28±0.75 | 0.60±0.27 |
| | *Capitellidae* spp. | 12.87±2.95 | 2.09±0.50 |
| | *Chaetozone* sp. | 13.29±1.99 | 5.39±1.27 |
| | *Cistenides hyperborea* | 15.92±9.15 | 0.27±0.12 |
| | *Cossura pygodactylata* | 465.73±64.87 | 28.56±6.88 |
| | *Dipolydora caulleryi* | 0.49±0.49 | 0.00±0.00 |
| | *Dorvilleidae* spp. | 30.12±6.07 | 1.82±1.46 |
| | *Dipolydora socialis* | 0.00±0.00 | 0.33±0.14 |
| | *Dysponetus pygmaeus* | 0.73±0.22 | 0.00±0.00 |
| | *Enipo canadensis* | 0.18±0.13 | 0.42±0.20 |
| | *Eteone flava* | 0.00±0.00 | 1.02±0.34 |
| | *Eteone longa* | 2.99±0.58 | 0.66±0.20 |
| | *Euchone incolor* | 3.29±1.71 | 0.72±0.40 |
| | *Goniada maculata* | 0.24±0.15 | 0.33±0.10 |
| | *Gyptis bruneli* | 16.40±1.66 | 2.24±0.73 |
| | *Lumbrineridae* spp. | 19.82±4.93 | 16.86±2.46 |
| | *Lysilla loveni* | 1.65±0.43 | 0.12±0.07 |
| | *Maldane glebifex* | 1.52±0.94 | 0.00±0.00 |
| | *Maldane sarsi* | 0.00±0.00 | 1.89±0.81 |
| | *Maldane* sp. A | 0.30±0.20 | 0.00±0.00 |
| | *Micronephtys neotana* | 18.41±1.93 | 6.04±0.81 |

| | | | |
|---|---|---|---|
| | *Nephtys ciliata* | 0.31±0.13 | 1.50±0.29 |
| | *Nereimyra aphroditoides* | 8.96±7.04 | 0.36±0.16 |
| | *Paradoneis lyra* | 0.00±0.00 | 0.99±0.45 |
| | Paraonidae sp. A | 0.55±0.38 | 0.00±0.00 |
| | *Pherusa plumosa* | 0.49±0.24 | 0.15±0.08 |
| | *Pholoe longa* | 0.55±0.31 | 3.07±1.52 |
| | *Pholoe minuta* | 0.96±0.10 | 0.96±0.41 |
| | *Prionospio steenstrupi* | 41.46±8.52 | 17.50±3.11 |
| | *Scalibregma inflatum* | 0.37±0.19 | 0.03±0.03 |
| | *Scoloplos armiger* | 1.77±0.99 | 2.22±0.71 |
| | *Sphaerodoridium minutum* | 1.65±0.63 | 0.18±0.08 |
| | *Spiochaetopterus typicus* | 0.12±0.09 | 0.42±0.15 |
| | *Syllides* sp. | 2.20±0.85 | 0.00±0.00 |
| | *Terebellides stroemii* | 3.35±0.68 | 0.57±0.16 |
| | *Terebellidae* sp. B | 0.85±0.40 | 0.00±0.00 |
| Other | *Antalis entalis* | 2.99±0.87 | 6.15±1.53 |
| | Chaetognatha | 0.12±0.09 | 0.18±0.07 |
| | *Ctenodiscus crispatus* | 0.18±0.10 | 0.45±0.13 |
| | Cumacean spp. | 5.24±1.45 | 3.01±0.80 |
| | Echinoidea | 0.55±0.21 | 0.06±0.04 |
| | Nemertea spp. | 10.49±1.76 | 4.80±1.25 |
| | Ophiuridae | 2.74±0.60 | 1.27±0.40 |
| | *Priapulus caudatus* | 0.00±0.00 | 0.30±0.17 |
| | Sipunculidea sp. | 2.44±0.56 | 0.48±0.15 |
| | Tanaidacea | 0.31±0.20 | 0.03±0.03 |

Note: to take a conservative approach, the following were grouped when considering "unique" species/taxa to historical and contemporary communities 1. *Maldane sarsi* and *Maldane* sp. A grouped as "*Maldane* sp." And 2. *Paradoneis lyra* and Paraonidae sp. A grouped as "Paraonidae sp."

**Table S3.** Abundance (total no. individuals) of species/taxa unique to historical and contemporary samples for major taxonomic groups (i.e., Amphipoda, Bivalvia, Gastropoda, Polychaeta, other).

| **Community** | **Species/Taxa** | **Major Group** | **Total abundance** |
|---|---|---|---|
| Historical | Ampharetidae sp. | Polychaeta | 38 |
| | *Amharete finmarchica* | Polychaeta | 10 |
| | *Dysponetus pygmaeus* | Polychaeta | 30 |
| | *Maldane glebifex* | Polychaeta | 63 |
| | *Dipolydora caulleryi* | Polychaeta | 20 |
| | *Syllides* sp. | Polychaeta | 90 |
| | *Terebellida* sp. B | Polychaeta | 35 |
| | *Artacama proboscidea* | Polychaeta | 73 |
| | *Yoldia* sp. | Bivalvia | 100 |
| | *Turridae* sp. | Gastropoda | 45 |
| | *Protomedeia* sp. | Amphipoda | 30 |
| | *Melita* sp. | Amphipoda | 18 |
| | *Quasimelita formosa* | Amphipoda | 13 |
| | *Bathymedon* sp. | Amphipoda | 58 |
| Contemporary | *Axinopsida orbiculate* | Bivalvia | 71 |
| | *Curtitoma incisula* | Gastropoda | 39 |
| | *Dipolydora socialis* | Polychaeta | 12 |
| | *Eteone flava* | Polychaeta | 37 |
| | *Megamoera dentata* | Amphipoda | 10 |
| | *Megayoldia thraciaeformis* | Bivalvia | 59 |
| | *Orchomenella minute* | Amphipoda | 27 |
| | *Paratryphosites abyssi* | Amphipoda | 201 |
| | *Priapulus caudatus* | other | 11 |
| | *Propebela rugulata* | Gastropoda | 23 |
| | *Trachyrhynchus erosus* | Gastropoda | 44 |

**Table S4.** Results of SIMPER analyses showing trait modalities contributing to ~50% of the dissimilarity between historical and contemporary communities for Placentia Bay, Newfoundland ordered from highest to lowest contribution. The cumulative percent of total expressed traits for influential modalities are also shown.

| **Modality (trait)** | **Dissimilarity (%)** | **Cumulative (%)** |
|---|---|---|
| High (tolerance) | 5.9 | 5.9 |
| Small (body size) | 5.7 | 11.6 |
| Subsurface deposit (feeding mode) | 5.6 | 17.2 |
| Benthic/direct (larval development) | 5.4 | 22.6 |
| Surface deposit (bioturbator) | 4.9 | 27.5 |
| Sexual external (reproduction) | 4.7 | 32.2 |
| Burrower (adult movement) | 4.7 | 36.9 |
| Burrowing (living habitat) | 2.6 | 39.5 |
| Crawler (adult movement) | 2.6 | 42.1 |
| Medium (body size) | 2.6 | 44.7 |
| Medium-large (body-size) | 2.5 | 47.2 |

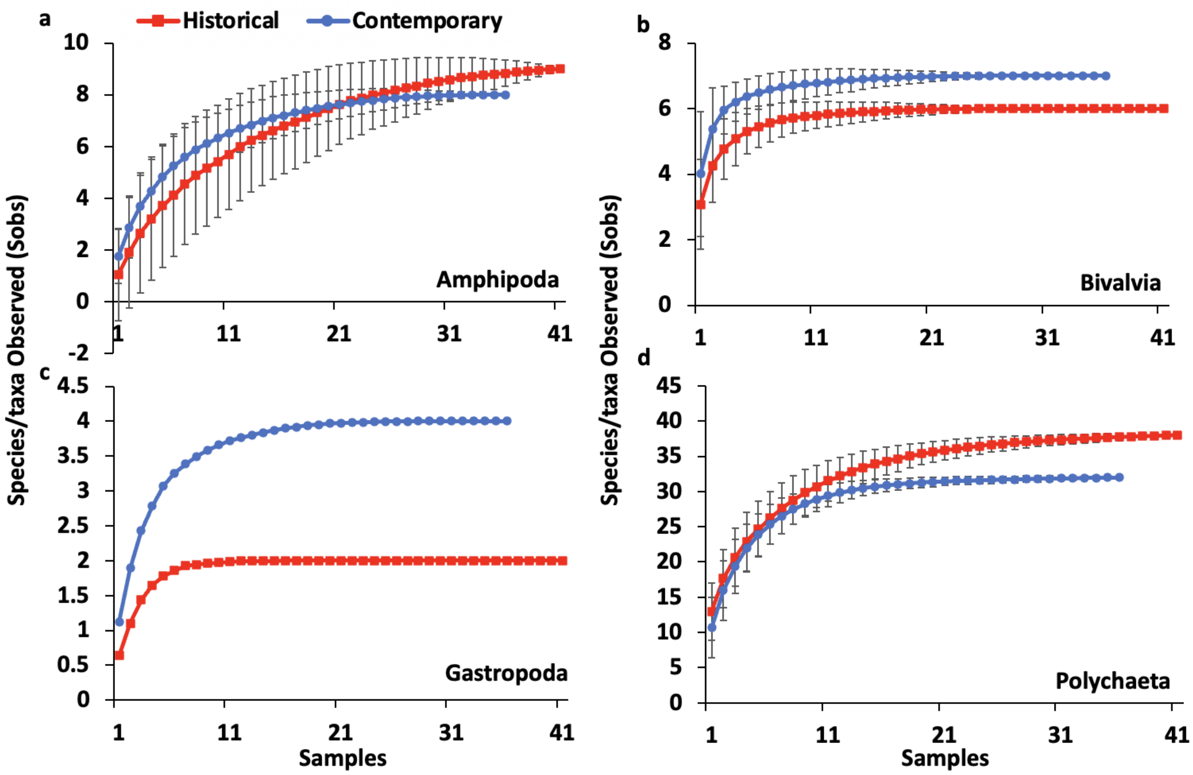


**Fig. S1.** Species accumulation curve based on species/taxa observations ±SD for major groups including (a) Amphipoda, (b) Bivalvia, (c) Gastropoda, and (d) Polychaeta for historical and contemporary samples.

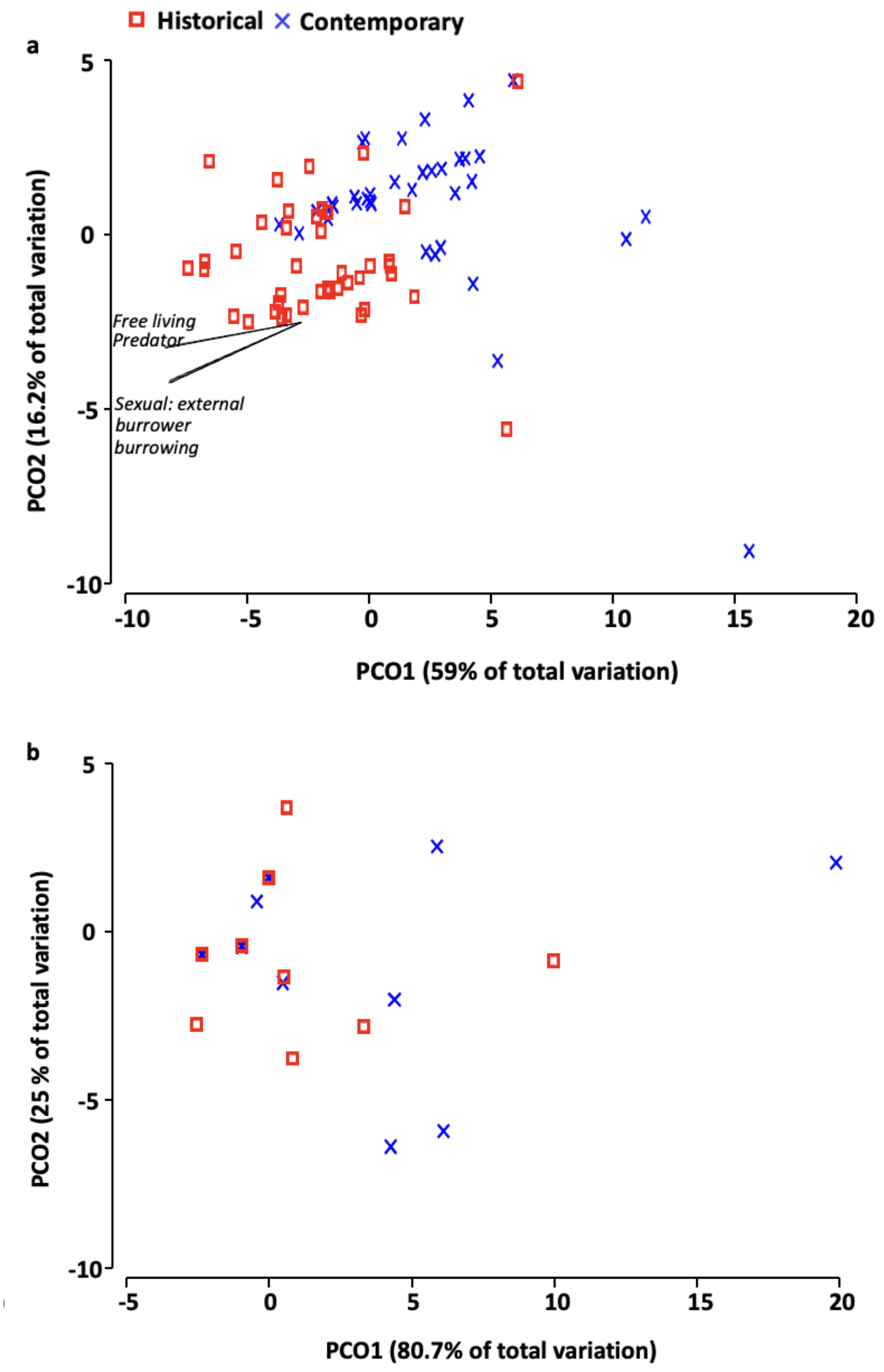


**Fig. S2.** Principal Coordinates Ordination (PCO) of functional traits for (a) expression of modalities based on Canberra similarity, and (b) presence/absence of modalities based on Sorensen resemblance. Differences between the historical versus contemporary communities were statistically significant (pseudo-$F_{1, 75}$= 19.5 and PERMANOVA P= 0.0001). Modality vectors=Pearson correlation of ≥ 0.7.

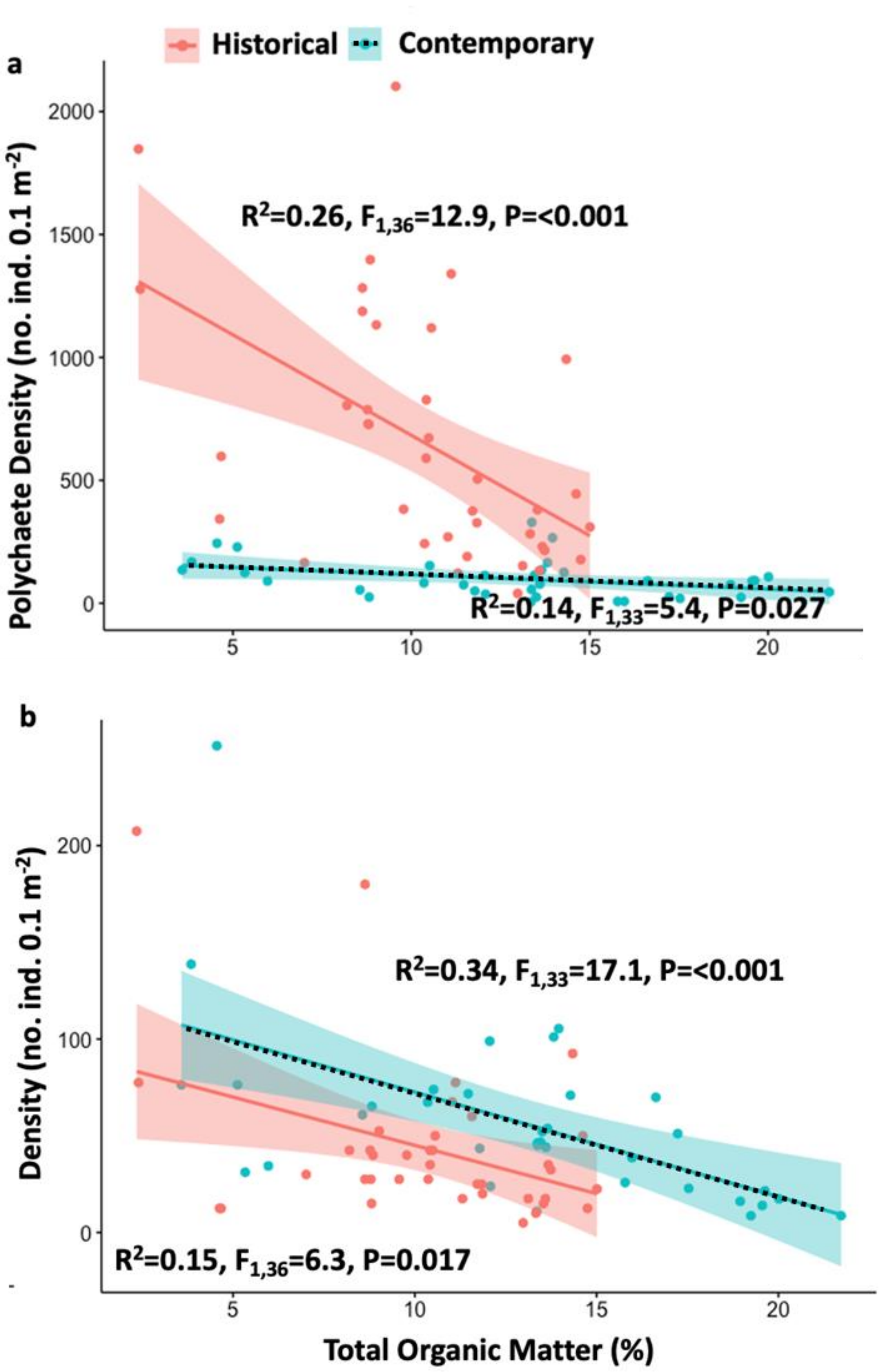


**Fig. S3**. Linear regression of density against total organic matter for (a) polychaetes (no. individuals $0.1m^{-2}$ ) and (b) major taxonomic groups excluding polychaetes (i.e., Amphipoda, Gastropoda, and Bivalvia combined). Historical=solid line and contemporary=dotted line.
